# Direct RNA sequencing enables improved transcriptome assessment and tracking of RNA modifications for medical applications

**DOI:** 10.1101/2024.07.25.605188

**Authors:** Charlotte Hewel, Anna Wierczeiko, Johannes Miedema, Felix Hofmann, Stephan Weißbach, Vincent Dietrich, Johannes Friedrich, Laura Holthöfer, Verena Haug, Stefan Mündnich, Lukas Schartel, Lioba Lehmann, Kristi Jenson, Stefan Diederich, Stanislav Sys, Tamer Butto, Norbert W Paul, Jonas Koch, Frank Lyko, Florian Kraft, Susann Schweiger, Edward A Lemke, Mark Helm, Matthias Linke, Susanne Gerber

## Abstract

Direct RNA sequencing (DRS) is a nanopore-based technique for analyzing RNA in its native form, promising breakthroughs in diagnostics and biomarker development. Coupled to RNA002 sequencing chemistry, its clinical implementation has been challenging due to low throughput, low accuracy, and lack of large-scale RNA-modification models. In this study, we evaluate the improvements achieved by pairing the latest RNA004 chemistry with novel modified-base-calling models for pseudouridine and *N*^6^-methyladenosine using diverse RNA samples from cell lines, synthetic oligos, and human blood. Finally, we present the first clinical application of DRS by confirming the loss of RNA methylation in a patient carrying truncating mutations in the methyltransferase *METTL5*. Conclusively, the combined use of RNA004 chemistry with the base-calling models significantly improved the throughput, accuracy, and site-specific detection of modifications. From this perspective, we offer an outlook on the potential suitability of DRS for use in routine diagnostics, the first comprehensive benchmark of human peripheral blood and quality assessments of RNA therapeutics.

## Introduction

Naturally occurring modifications to RNA such as N^6^-methyladenosine (m^6^A) and pseudouridine (Ψ) crucially affect its structure, stability, and its interactions with proteins, and as such dynamically regulate molecular processes in cells. More than 170 chemical RNA modifications are currently known, and more are expected to be discovered(1).

Modifications on mRNA molecules appear to be involved in translating, splicing, and stabilizing RNA(2, 3). For example, pseudouridylation at stop codons can enable readthrough, allowing protein synthesis despite a “translation to stop” signal (4). This mechanism has attracted significant attention in drug development, as approximately 10–20% of genetic mutations reported in the variant database ClinVar contain premature termination codons (PTCs). PTCs give rise to truncated proteins that cannot function as intended, leading to various inherited diseases. Translational readthrough-inducing drugs (TRIDs) show promise as therapeutic agents for a number of rare diseases(5). Recent advances in this field include work by Schartel et al., who developed a model organelle system using DKC1 and small nucleolar RNAs (snoRNAs) as guide RNAs to achieve precise, site-specific pseudouridylation, enabling controlled translational readthrough at targeted transcripts(6).

Several aberrations of RNA-modifying enzymes are linked to human diseases, so called “modopathies” (7–9). For example, loss of pseudouridine synthase PUS1 is associated with mitochondrial myopathy with lactic acidosis and sideroblastic anemia (MLASA), whereas dysfunctional PUS3 and PUS7 are associated with intellectual disability and neurodevelopmental delay (10–12). Additionally, patients with dyskeratosis congenita have reduced pseudouridylation of 28S rRNA or the telomerase RNA component (TERC) (13, 14).

m^6^A plays an important role in multiple cancers (15). For example, the methyltransferase METTL3 can be upregulated in glioblastoma, thereby upregulating the expression of the target cancer gene SOX2. On a related note, epitranscriptomic rRNA fingerprinting approaches, to distinguish between tumor and normal samples from a fraction of reads - or ultra low depth sequencing- show promising avenues to classify cancers based on their epitranscriptomic signature at either a fraction of the cost or a fraction of the time classical approaches would take (16).

Interestingly, the methyltransferase METTL5 is known to be responsible for the methylation of a specific adenine at position 1832 of the 18S rRNA, and is therefore essential for the translation process (17). Dysfunction of METTL5 due to genetic mutations causes an intellectual developmental syndrome with severe microcephaly (18).

Consequently, both human diagnostics and research applications would benefit from a streamlined and high-throughput method for measuring RNA modifications.

Conventional RNA sequencing (RNA-seq) by next-generation sequencing (NGS) facilitates differential expression analysis of genes or transcripts and analysis of differential splicing in a high-throughput mode (19). However, the required conversion from RNA into cDNA and the subsequent fragmentation erases a sizable quantity of information present on native RNA, such as modifications. Thus, conventional RNA-seq is not suited to directly observe either full-length isoforms or RNA modifications (20). For clinical and research applications this indicates that metrics for RNA modifications could only be provided by indirect measurements, such as differences in the level of methyltransferase expression or splice isoforms.

Direct RNA sequencing (DRS) by Oxford Nanopore Technologies (ONT) is a major innovation, as it enables the detection of nucleotide-specific modifications directly on native RNA molecules by measuring real-time variations in electrical current (21). DRS can quantify gene expression while simultaneously capturing full-length transcripts, splicing patterns, poly(A)-tail length, and distinct RNA modifications within a single assay (22), highlighted are the use cases that are covered within this manuscript (Fig 1).

**Figure 1.**
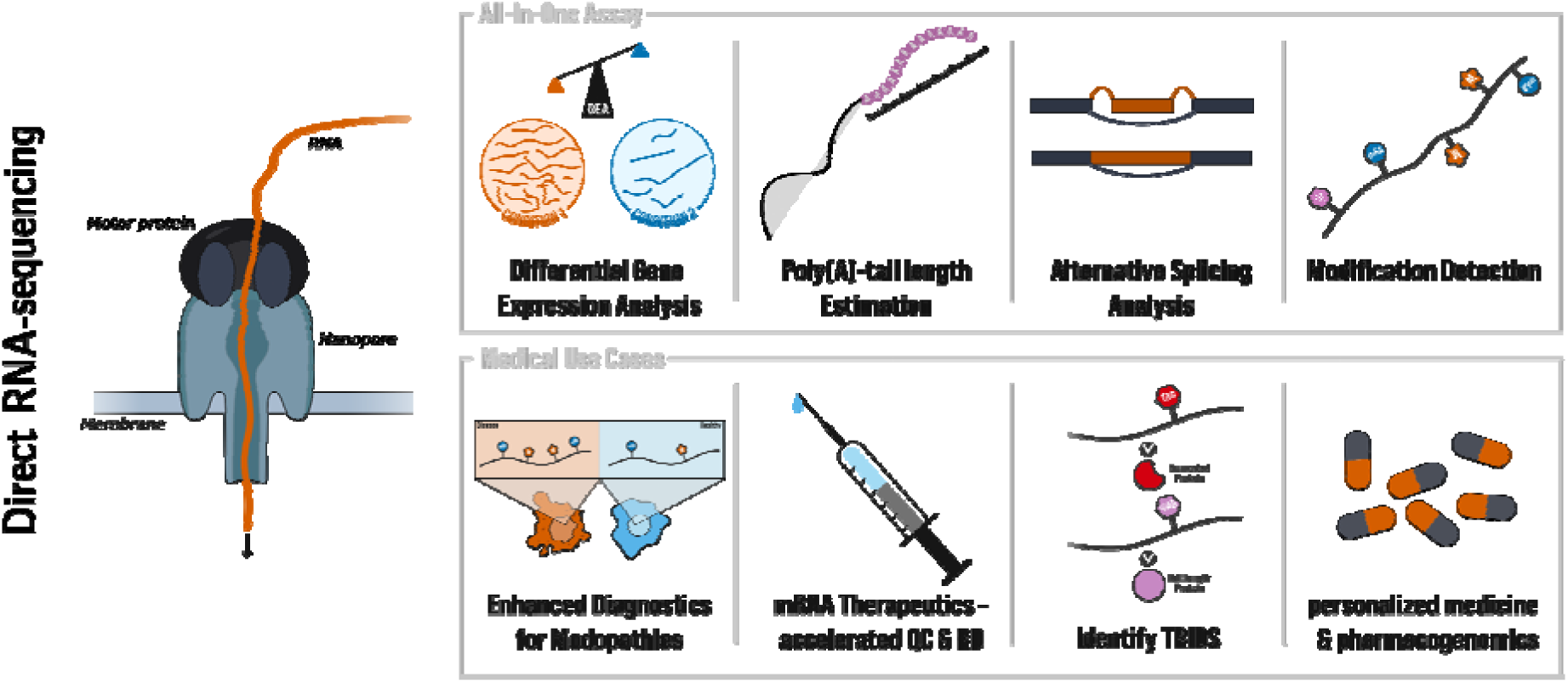
Selected uses for DRS in clinical therapy and analysis options. In addition to the analysis of differential gene expression and alternative splicing, DRS can be utilized to estimate poly(A)-tail lengths and to predict Ψ and m6A modification sites. These features provided by the latest base-calling model enable the integration of DRS in a clinical context, such as diagnostics, the development of RNA therapeutics, and identification of TRIDS. TRID, translational readthrough inducing drug; RD, research and development; QC, quality control.The shaded boxes represent the subset of use cases addressed in this manuscript.

This capability for comprehensive profiling holds promise for advanced diagnostic and research applications, including enhanced detection of modopathies, accelerated development and quality control of mRNA therapeutics, and simplified epitranscriptomics analyses (23, 24)

Although these prospects are promising, it is important to acknowledge the evolution of DRS technology. The chemistry of the now discontinued SQK-RNA002 sequencing kit for DRS was applied in various contexts, such as tRNA sequencing (25). However, its performance was mixed owing to its low throughput, low accuracy, and the absence of modified-base calling within ONT basecallers (26, 27). To address these limitations, ONT introduced the SQK-RNA004 sequencing kit, featuring updated flow cells, a new motor protein, and base-calling models capable of detecting the RNA modifications m^6^A, Ψ and canonical nucleotides.

In this study, we comprehensively compare the RNA002 and RNA004 chemistries using diverse RNA samples including cell cultures and human blood, evaluating improvements in yield, quality, gene coverage, poly(A)-tail length estimation, and the detection of m^6^A and Ψ modifications. To illustrate the practical implications of using RNA004 chemistry, we highlight two practical applications of DRS, a model system for future RNA therapeutics and the diagnostic of a rare disease case.

First, we present an example for the performance of site-specific Ψ detection in RNA therapeutics by validating the expected stoichiometry introduced by a pseudouridylation system developed by Schartel et al. 2024, using both the RNA002 and RNA004 flow cells(6).

Second, we showcase the first clinical application of direct RNA sequencing using RNA004 chemistry. Here, we confirmed the loss of function of the m^6^A methyltransferase METTL5 in a patient harboring two compound heterozygous variants predicted to disrupt enzyme activity, classified as pathogenic and of uncertain significance (VUS). This highlights how DRS can improve the interpretation of VUS in RNA-modifying enzymes and be a promising tool for clinical diagnostics.

## Material and Methods

### Sample description

Five different sample sources were used during this study and sequenced a total of 21 times. Universal Human Reference RNA (UHRR) was purchased from Thermo Fisher Scientific (cat. no. QS0639). The HEK293T cells were transfected with EGFP and mCherry. The human samples were taken from healthy volunteers or a patient after written informed consent was obtained.

### HEK293T samples

HEK293T cells were transfected with artificial snoRNAs as well as EGFP and mCherry. The snoRNAs were designed to target a premature stop codon within the EGFP and mCherry transcripts at nucleotide positions 115 and 565, respectively. In Case A, both mRNAs were targeted for pseudouridinylation; in Case B, the mCherry mRNA was preferentially targeted owing to a decrease in EGFP pseudouridinylation. In the Control condition, a scrambled snoRNA was transfected, as were mCherry and EGFP (6).

### Direct RNA library preparation for the cell line samples and RNA002/RNA004 chemistries

For direct RNA library preparation, we used either the old DRS chemistry (SQK-RNA002, ONT) or the updated kit (SQK-RNA004, ONT) following the manufacturer’s protocol. In brief, 100 ng of poly(A)-tailed RNA or 1000 ng of total RNA was adjusted to 9 μl with nuclease-free water. To this RNA sample, 3 μl of NEBNext Quick Ligation Reaction Buffer (New England Biolabs, B6058), 1 μl RT Adapter (RTA, ONT), and 1.5 μl T4 DNA Ligase (2×10^6^ U/ml; New England Biolabs, M0202) were added, resulting in a total volume of 14, 5 μl. The reaction was mixed by pipetting and incubated for 10 min at room temperature. Next, the reverse transcription master mix was prepared by mixing 9 μl of nuclease-free water, 2 μl of 10 mM dNTPs, 8 μl of 5× first-strand buffer (Thermo Fisher Scientific), and 4 μl of 0.1 M DTT. This master mix was added to the RNA sample containing the RT Adapter-ligated RNA along with 2 μl of SuperScript III reverse transcriptase. The reaction was incubated at 50°C for 50 min then at 70°C for 10 min, and then cooled to 4°C. RNAClean XP beads (72 μl; Beckman Coulter, A63987) were then added to the reaction, followed by incubation on a Hula mixer for 5 min at room temperature. Subsequently, the sample was washed twice with 70% ethanol, and the DNA was eluted with 20 μl of nuclease-free water. The eluted DNA was used in the adapter ligation reaction. For that reaction, 8 μl of NEBNext Quick Ligation Reaction Buffer, 6 μl of RNA Adapter (RMX for RNA002; RLA for RNA004), 3 μl of nuclease-free water, and 3 μl of T4 DNA Ligase were mixed with the 20 μl of eluted DNA (total volume: 40 μl). The reaction was incubated for 10 min at room temperature. After incubation, 20 μl of RNAClean XP beads were added to the adapter ligation reaction, followed by incubation on a Hula mixer for 5 min at room temperature. The sample was then washed twice with Wash Buffer (WSB, ONT) using a magnetic rack. Next, the pellet was resuspended in 41 μl (RNA002) or 33 μl (RNA004) of Elution Buffer (EB) and incubated at 37°C for 10 min in a Hula mixer to release long fragments from the beads. Finally, the eluate was cleared by pelleting the beads on a magnet, retained, and transferred to a clean 1.5 ml tube. One microliter of reverse-transcribed and adapted RNA was quantified using a Qubit fluorometer. For R9.4.1 PromethION sequencing (RNA002), 40 μl of the library was mixed with 35 μl of nuclease-free water and 75 μl of RRB and loaded into a R9.4.1 PromethION flow cell. For PromethION sequencing (RNA004), 32 μl of library was mixed with 100 μl of Sequencing Buffer (SB) and 68 μl of Library Solution (LIS) and loaded into an RNA chemistry PromethION flow cell. For the 18S rRNA sample, a MinION RNA flow cell (FLO-MIN004RA) was loaded in accordance with the manufacturer’s instructions.

### Peripheral blood and *in vitro* transcription

The peripheral blood was obtained from a healthy volunteer. The RNA was extracted using the PAXgene Blood miRNA Kit from Qiagen according to the manufacturer’s protocol, except the RNA was eluted in nuclease-free water instead of the buffer provided. The RNA was characterized using the Bioanalyzer total RNA Nano Assay according to the manufacturer’s protocol. Depletion of globin mRNA was performed with the GLOBINclear-Human Kit from Thermofisher Scientific (AM1980) according to the manufacturer’s protocol; this was carried out four times. The total input of RNA was 20 μg, the total output was 11 μg of globin-depleted RNA. The concentration was measured using the Qubit RNA HS Assay from Thermofisher Scientific. Five micrograms of RNA was stored for later use in the direct RNA Run. The subsequent reverse transcription (RT), PCR, IVT and polyadenylation were carried out according to Tavakoli et al. (2023). The following individual amendments were made: the IVT primers used in the PCR had a final concentration in the reaction of 1 μM per primer. The input amount of RNA used for the RT and PCR was 632 ng; the output was 3910 ng of cDNA, measured with the Qubit DNA HS Assay (Thermo Fisher Scientific). The remaining RNA was degraded using the RNase Cocktail from Thermo Fischer Scientfic (AM2286) for 10 minutes at 37°C.The IVT was carried out with an input of 1140 ng template cDNA. The output was 40 μg RNA, as measured with the Qubit RNA HS assay. The remaining template cDNA was degraded using DNase I from the Paxgene miRNA Kit according to the manufacturer’s protocol. Libraries were prepared using the SQK-RNA004 sequencing kit (ONT). RNA input for the IVT was 4, 5µg and for the direct RNA runs 5µg. The library output was 295 ng of RNA/cDNA hybrid for the IVT and 700ng for the direct RNA runs, as measured with the Qubit DNA HS Assay. The librarys were loaded completely onto PromethION RNA Flow Cells (FLO-PRO004RA).

### Base calling and alignment of RNA002 and RNA004 runs

The raw pod5 files from all RNA004 sequencing runs were base-called using Dorado v0.7.2 with the canonical base-call model <u>rna004_130bps_sup@v5.0.0</u>. The model allowed for direct calling of m^6^A and Ψ using the flag --modified-bases m6A pseU. poly(A)-tail lengths were also estimated by including flag --estimate-poly-a, after the tailfindr algorithm that was recently adopted by ONT. The base calling of raw pod5 files from the RNA002 sequencing runs was done with Dorado’s high accuracy model for RNA002, that is, rna002_70bps_hac@v3. Base-called reads of all samples were then aligned to the primary assembly of the human reference genome hg38, downloaded from Gencode release 43 (https://ftp.ebi.ac.uk/pub/databases/gencode/Gencode_human/release_43/GRCh38.primary_assembly.genome.fa.gz). In addition, the model from Diensthuber et al., was used on the RNA002 samples for benchmarking purposes (28). Alignment was performed in Minimap2 v2.26 with the following settings: -y --MD -ax splice -uf -k14. The resulting BAM files were sorted and indexed using samtools v1.16.1. The HEK293T samples were additionally mapped onto the EGFP and mCherry reference sequences for analyzing the detection of modified targets. The oligos were mapped in addition to their custom oligo references (see Schartel et al. 2024). The quality metrics of all sequencing runs and mappings were derived by NanoComp v1.23.1. The average base-call quality and the alignment-based percent identity were visualized in Python 3.8 using matplotlib v.3.8.3 and seaborn v.0.13.2. The percentage mismatch on chromosome 20 for the cell line samples was performed using dRNA-eval, after realignment as described on GitHub (https://github.com/KleistLab/nanopore_dRNAseq)(29) and subsequently plotted in python.

### Annotation of genomic features

Reads were mapped to genomic features using featureCounts v.2.0.6. Basic gene annotation downloaded from Gencode v43 served as the annotation reference (https://ftp.ebi.ac.uk/pub/databases/gencode/Gencode_human/release_43/gencode.v43.basic.annotation.gtf.gz). Parameter –L was passed to featureCounts to account for long reads as input; –s 0 to perform unstranded read counting. The format of the annotation file was specified with -F ’GTF’.

### Transcriptome and Mendeliome counts

The feature counts per sample generated by featureCounts v2.0.0 were normalized by using DESeq and a Principal Component Analysis (PCA) was done using the top 500 most variable expressed genes across samples. The PC1 and PC2 were plotted using python. Furthermore, those genes with at least 10x coverage were intersected with an in-house list of genes associated with known diseases (Mendeliome). Both the number of total genes and Mendeliome genes with coverage >= 10 reads were visualized for all samples using python’s seaborn and matplotlib libraries.

### Modification information extraction

The modification bed files were generated from the Dorado base-called modbam files with modkit version 0.3.1. For m^6^A the reads were subset to DRACH regions with the flag –motif DRACH 2, additionally the flags --ignore 17802 and --filter-threshold A:0.8 -- mod-threshold a:0.98 were used, as determined by the modification probability histogram also made with modkit. For Ψ the flags --ignore a, --filter-threshold T:0.8, and -- mod-threshold 17802:0.98 were used. Then the bedfiles were filtered to have a valid coverage of at least 10 reads and a site-specific methylation of at least 10% to reduce false positives. This cutoff was defined by calculating precision and recall for the blood m6A samples based on the difference between sites in the m6A dorado-set and the orthogonal GLORI set at different coverage cutoffs.

### m6A and pseU analysis

For both m6A and pseu sites with coverage >= 10 reads and m6A ratio >= 10 %, all pairwise correlations of m6A ratios from intersected sites were done by calculated the Pearsons correlation coefficient using pearsonr from scipy.stats in python. For sample replicates, the number of overlapping sites were plotted as venn diagrams using matplotlib_venn in python. We both plotted the overlap of all detected sites as well as those sites that were covered with at least 10 reads in all replicates, respectively. The read depths for all modification sites and samples were retrieved by using samtools depth with default options on both modification bed files and aligned bam files. For RNA004 blood samples, we additionally plotted the m6A ratios per sample for all sites detected in each sample as well as for joined covered sites.

### Nanopore methylation control data

*The raw data for the modified and unmodified control oligos for m6A and pseudouridine, that ONT used for model development and benchmarking were downloaded from the following AWS instance: s3://ont-open-data/rnamodbase-validation_2025.03. And a detailed description of the dataset can be found here:* https://epi2me.nanoporetech.com/rna-mod-validation-data/*. Subsequently, the data was rebasecalled to ensure usage of consistent basecaller settings*.

### Analysis of **Ψ** and m^6^A detection at target positions

For the site-specific analysis of modification, the Dorado-derived modification probabilities as well as the mismatch frequencies were extracted from the ML/MM tags of the respective bam files using pysam v0.22.1 with min_base_quality=13 and threshold=0.98 (Python 3.8, pysam: https://github.com/pysam-developers/pysam). Additionally, we performed Dorado-based Ψ calling on reads harboring misbasecalled Cs by changing the motif specification to motif=“C“ in the conFiguretoml file of the respective Dorado base-calling model.

For the EGFP and mCherry motifs as well as the known Ψ-site on the PSMB2 transcript, the frequencies of U-based Ψs, C-based Ψs, unmodified Cs and unmodified Us were calculated and plotted in R v4.2.2 using the R package ggplot2 v3.4.4.

To extract the m6A modification frequencies for all 18S rRNA transcripts, we additionally mapped the raw reads onto the rDNA reference sequence published by George et al.(30) for the peripheral blood samples from two healthy individuals and one patient, as well as the 18S rRNA IVT sample. The m6A modification probabilities at the 18S rRNA position A1832 for the peripheral blood and the 18S rRNA IVT samples were extracted using pysam with min_base_quality=13 and both the probability distribution and the m6A stoichiometry of modified sites with an probability threshold>=0.98 at the A1832 site were plotted using the Python package seaborn v0.13.2.

### Estimation of poly(A)-tail length

Poly(A) features were extracted during base calling with Dorado by adding the flag –estimate-polya as detailed above. For the basic comparison between tailfindr and Dorado 0.7.2 a test data set for RNA002 chemistry was downloaded from ERR3349888. The raw single fast5 data was subsequently transferred into multi fast5 via single_to_multi_fast5 from ont-fast5-api toolkit and transferred into pod5 via pod5 convert fast5. Then base calling was performed with Dorado 0.7.2 rna002_70bps_hac@v3 model and --estimate-poly-a flag. The poly(A) length was extracted from the resulting ubam file by storing the pt tag of each read in a table. For the tailfindr length estimation, we made use of the information on pre-existing length and the barcoding table as provided by the analysis of Krause and Niazi (https://github.com/adnaniazi/krauseNiazi2019Analyses). Subsequently, the data was loaded in Jupyter Notebook and plotted with seaborn. For the RCS poly(A) reads aligning to the RCS reference were extracted from the first run of UHRR on RNA002 and RNA004 respectively and subsequently plotted. The expected tail length of this oligo is around 30 bp. The poly(A)-tail length distribution of all aligned reads was plotted per sample. Additionally, we calculated the mean poly(A)-tail length per gene by extracting the pt tags from all primary mapped reads per gene region using pysam. The mean poly(A)-tail lengths distribution of genes with minimum coverage of 10x was plotted per sample and the pairwise correlation of mean poly(A)-tail lengths from covered genes was calculated and plotted between replicates. Furthermore, we extracted the pt tag for those reads and mapped on the genes *DDX17*, *OLA1* and *SRP14* for RNA002 and RNA004 HEK293T samples and plotted the poly(A)-tail length distribution per sample and gene in seaborn, specifically.

### m^6^A calling using mAFiA and m6ABasecaller for chromosome 20

The data was subset to chromosome 20 via filtering by samtools. Then, pod5 filter was used on the read IDs to retain a subset of the raw data for chr20. pod5 convert to_fast5 was used to transfer data into fast5 as required for downstream analysis with the base callers for m^6^A RNA002. Both mAFiA and m6Abasecaller were run with default options as described in (https://github.com/dieterich-lab/mAFiA & https://github.com/novoalab/m6ABasecaller). The GLORI test data set was obtained from Liu et al. (2023)(31). The GLORI set for the peripheral blood was derived from two replicates, that were sequenced in house. Dorado 0.7.2 and modkit were run as described previously. Plotting was done in Python using UpSetPlot version 0.9.0.

### Coverage subsampling for RNA004 vs RNA002

In order to evaluate how many sites would have been called if the RNA004 blood samples had equal throughput to the RNA002 samples. The reads in all files were counted via samtools and subsequently the coverage was of the RNA004 files was downsampled to the fraction of coverage from the RNA002 sample via samtools. Then, the modkit step was repeated as previously described.

### RNA isolation and preparation for GLORI and direct RNA control sequencing

Total RNA from 3 biological HEK293T replicates was isolated using TRIzol. Small RNA species were depleted using the MEGAclear Transcription Clean-Up Kit (Thermo Fisher Scientific). mRNA enrichment was performed twice using the Dynabeads mRNA Purification Kit (Thermo Fisher Scientific). The same step was repeated for two biological replicates of the blood of the healthy control person.

For direct RNA sequencing of the HEK293T cells, 300 ng of mRNA pooled from the three biological replicates were sequenced on a single flow cell on the MinION Mk1B platform using the direct RNA sequencing kit (SQK-RNA004; Oxford Nanopore). Data analysis was performed as detailed above.

For GLORI sequencing, the mRNA was fragmented at 94 °C for 3 min using the NEBNext Magnesium RNA Fragmentation Module (New England Biolabs) and purified using the RNA Clean & Concentrator-5 kit (Zymo Research) including a DNase I-digestion step. mRNA protection, deamination, and deprotection were performed as described in literature (31, 32). For preparing the sequencing libraries, RNA samples were end-repaired via Antarctic phosphatase (New England Biolabs) and T4 Polynucleotide Kinase (New England Biolabs) treatments according to the manufacturer’s instructions. End-repaired samples were purified using the RNA Clean & Concentrator-5 kit (Zymo Research). Sequencing libraries were then prepared using the NEBNext Small RNA Library Prep Set for Illumina in combination with the NEBNext Multiplex Oligos for Illumina (Index Primer Sets 1 and 3) (New England Biolabs). Sequencing was performed by the Next Generation Sequencing Core Facility of the German Cancer Research Center, Heidelberg on a NovaSeq 6000 platform (Illumina) using a 100 bp pairedend sequencing protocol. Sequencing adaptors from raw reads were removed by Trim Galore (version 0.6.6). Trimmed reads were further processed by the GLORI-tools pipeline. GLORI-tools is available on GitHub: https://github.com/liucongcas/GLORI-tools. Software used for executing the GLORI-tools pipeline included python (version 3.10.1), samtools (version 1.19), STAR (version 2.7.10a), and bowtie (version 1.3.0). The human genome (GRCh38) and transcriptome (GCF_000001405.39) reference files were obtained from UCSC. To investigate the correlation of methylation ratios between DRS and GLORI-seq samples of HEK293T cells, replicates were merged by averaging the methylation ratios across overlapping m6A sites. Bivariate density plots were generated using ggplot version 3.5.1, the goodness-of-fit measure R^2^ was calculated using base R version 4.3.2 to assess the correlation between methylation ratios.

### Base-calling error pattern extraction and pseudouridine calling using NanoCEM in reporter sequences EGFP and mCherry

nanoCEM version 0.0.6.1 was run with default options for the HEK293T samples A, B, and C, and positions 115 and 565 for the sequences of EGFP and mCherry.

### U–C mismatch analysis on high-confidence pseudouridine sites

The U>C mismatch frequencies for known pseU sites detected by either Tavakoli et al. or listed in the RNA modification database RMBase(cite) were extracted from the bam files of all samples using pysam’s pileup function in python. The U>C mismatch frequency was calulcated by dividing the number of reads with U-calls by the total number of reads mapped onto the same position.

### DRACH-motif analysis

To determine the 5mer context of m6A sites, we extracted the +/-2 base flanking sequence from the GRCh38 reference genome for all sites with minimum coverage of 10 reads and an m6A ratio >= 10 % from the modkit bed file of each sample using Biopython’s SeqIO. We visualized the m6A ratio per DRACH motif as well as the proportion of each modified DRACH motif used per sample. Furthermore, we calculated the ratio of the mean m6A frequency observed in IVT and the mean m6A frequency observed in native RNA blood samples for each DRACH motif.

### 18S rRNA Methylation Control Sample Plasmid preparation and *in vitro* transcription

The target sequence was cloned into a pUC57 vector, which included an internal T7 promoter, the desired template sequence, and a BshTI restriction enzyme site at the 3’ end. Linearization of the plasmid was carried out overnight, following manufacturer’s instructions (Thermo Fisher Scientific). Next, the plasmids were purified using phenol–chloroform extraction followed by ethanol precipitation. Successful linearization and the quality of the plasmids were confirmed by agarose gel electrophoresis and analysis with a NanoDrop One spectrophotometer.

IVT was carried out using the HiScribe T7 High Yield RNA Synthesis Kit (New England Biolabs) according to the manufacturer’s instructions. In brief, 2 µg of linearized plasmid was used as the template, along with 10× Reaction Buffer, 10 mM NTPs, and 2 U of T7 RNA Polymerase Mix. The reaction mixture was incubated at 37°C for 2 hours, and the process was stopped by digesting the template plasmid with DNase I (Thermo Fisher Scientific, EN0525) according to the manufacturer’s protocol. The resulting RNA was purified using the Monarch RNA Cleanup Kit (New England Biolabs, T2040), and the quality of the product was evaluated by capillary electrophoresis using Agilent RNA ScreenTape Analysis.

### Patient sample and data processing

Genomic DNA was isolated from the patient’s blood sample. Subsequently, all coding exons including flanking intron sequences of genes were enriched (“target enrichment” by hybridization) up to positions +/-20 using the SureSelect QXT Exome V7 Enrichment System (Agilent). The 2×150 bp (pairedend) NGS was performed on the NextSeq 500 System (Illumina) using the NextSeq 500/550 High-Output v2 Kit (300 cycles) reagents (Illumina).

The sequenced Illumina data were first converted to fastq files using bcl2fastq v2.20.0.422 and subsequently mapped onto the human reference genome hg19 using the bwa-mem aligner integrated in the Clara Parabricks Workflows (*pbrun fq2bam*) from NVIDIA (version 4.0.0-1).

#### *METTL5* RT-PCR assay

To determine splice aberration in the patient sample, a *METTL5*-gene-specific PCR was performed. The RNA was reverse-transcribed into cDNA using the PrimeScript RT Reagent (Takara) according to the manufacturer’s protocol. cDNA was amplified using a *METTL5* gene-specific PCR, targeting exons 1–7. Primer sequences are provided in Table S11. The FastStart High Fidelity PCR System (Roche) was used according to the manufacturer’s protocol, except that only a 25 µl reaction was prepared. The annealing temperature was 60°C. The elongation time was 2 min, with 35 cycles in total. The product was quantified using the Qubit DNA BR Assay.

Library preparation was performed using the Ligation Sequencing Kit (SQK-LSK114, ONT). The library was loaded completely into a PromethION DNA Flow Cell (FLO-PRO114M) and sequenced for approximately 9 hours.

The data was aligned against GRCh38 and only reads mapping to *METTL5* were retained by filtering with samtools. Subsequent plotting was done with ggsashimi v1.1.5.

## Results

### Enhanced performance and yield of RNA004 chemistry in direct RNA sequencing

A total of 25 flow cells using either RNA002- or RNA004-compatible settings were sequenced on a PromethION sequencer and two test samples on a MinION (Figure 2A, Table S1). First, we compared the performance of the two chemistries in terms of throughput and quality. To this end, we sequenced three different types of samples on a PromethION device using both RNA002 and RNA004 chemistry: 1) Universal Human Reference RNA (UHRR; total RNA), 2) poly-selected RNA from HEK293T cells, and 3) total RNA from human peripheral blood from a healthy individual (Figure 2A). All runs were base-called using the recently released Dorado base-caller (version 0.7.2, ONT). UHRR was used as a technical control to ensure comparability with previously published data from ONT. The HEK293T cells were additionally transfected with a newly developed pseudouridylation system and two mRNA reporter sequences (EGFP and mCherry) carrying Ψ-target sites, which were utilized later in the study to evaluate the detection of RNA modifications (Figure 2A, Table S1)(6). The RNA obtained from human blood was sequenced both in a native form and after *in vitro* transcription (IVT). IVT incorporates unmodified RNA nucleotides only and erases natural poly(A)-tails from the native molecules and is therefore used as an unmodified control for the purpose of detecting modifications (see Material and Methods, Figure 2A, Table S1).

**Figure 2.**
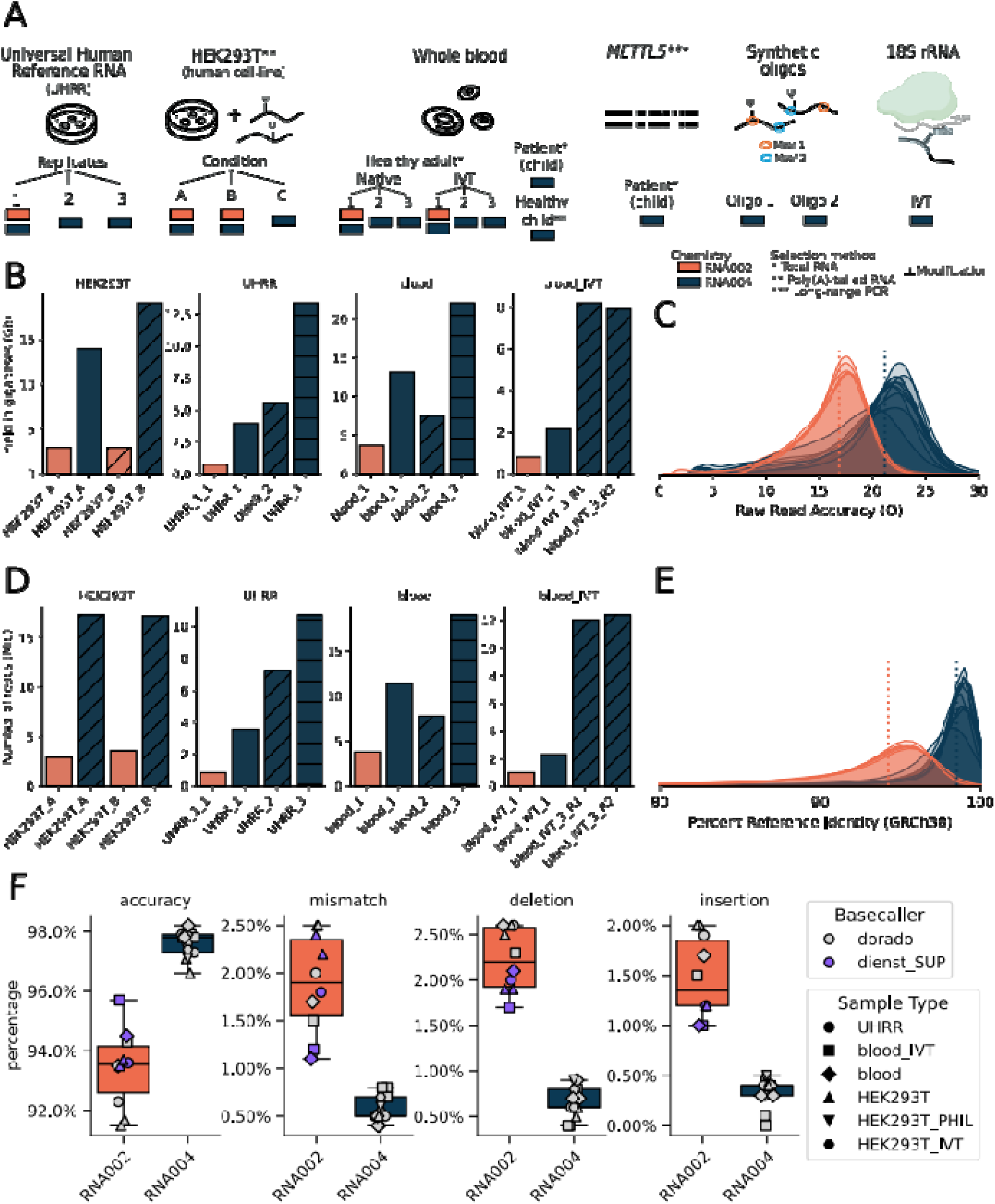
General sequencing quality metrics for all samples. Samples sequenced with the RNA002 chemistry and RNA004 chemistry are shown in orange and blue respectively. Samples which were basecalled using the SUP model provided by Diensthuber et al. (2024) are shown in purple. (A) Overview of sample types and number of replicates sequenced. The HEK293T cells contain the sequences of mCherry and EGFP, each with a premature stop codon targeted for pseudouridylation. (B) Yield in gigabases for the main human-derived samples (based on NanoComp), as well as the IVT samples derived from blood. (C) Raw read accuracies. (D) Number of reads for all samples (based on NanoComp). (E) Density plot of the percent reference identities for all samples as measured against GRCh38 (NanoComp), (F) For chromosome 20, the accuracy, mismatch deletion and insertion for both chemistries (RNA002 and RNA004) and different basecalling models was evaluated.

Following sequencing, we determined that the overall yield is dependent on the chemistry type (RNA002 or RNA004), method of library preparation (poly(A)-selected or total RNA), and sample origin (standardized cell line or peripheral blood; Figure 2B, D; Table S1). For all samples of the same composition, the RNA004 chemistry delivered higher yields than RNA002 (Figure 2B, D; Table S1).

The IVT samples, using either chemistry, showed the lowest throughput with < 2 gigabases (Gb) and < 3 million reads. Subsequent resequencing of a fresh batch with an improved protocol vastly improved throughput to < 25 Gb and < 30 million reads. The highest yield in Gb for a single flow cell overall was achieved by replicate 3 of the peripheral blood sample with < 30 Gb and < 22 M reads. The HEK293T cells using RNA004 chemistry, achieved a yield of approximately 17.3 and 21.94 Gb and more than 18 million reads. By contrast, the same samples sequenced with SQK-RNA002 yielded less than 35% of the throughput observed with RNA004 and < 7 million reads (Table S1).

Consistent with a general increase in throughput, the RNA004 runs show an increased average base quality derived from the Phred-based Q-scores, as well as better percent reference identity than the RNA002 chemistry, with mean scores close to 98% (Figure 2C, E, F; Table S1). For both chemistries, the most frequent source of base-calling errors were deletions, which is a common bias for ONT data(33, 34). As before, RNA004 chemistry outperforms RNA002 chemistry, even when using the fine-tuned model from Diensthuber for RNA002, which has higher accuracies than the ONT standard model for RNA002 (Figure 2F)(28).

### Enhanced gene expression profiling with RNA004

We further analysed the transcriptional patterns, including gene expression to assess the utility of RNA004 chemistry in clinical settings. From the general gene expression pattern, the samples separate chiefly by biotype and to a much lower extent by chemistry, as can be seen by the PCA plot in Figure 3A.The number of genes covered with at least 10 reads (10X) was lower if RNA002 chemistry was used (Figure 3B). This observation correlates with the higher throughput observed for the RNA004 chemistry (Table S1, Table S2).

**Figure 3.**
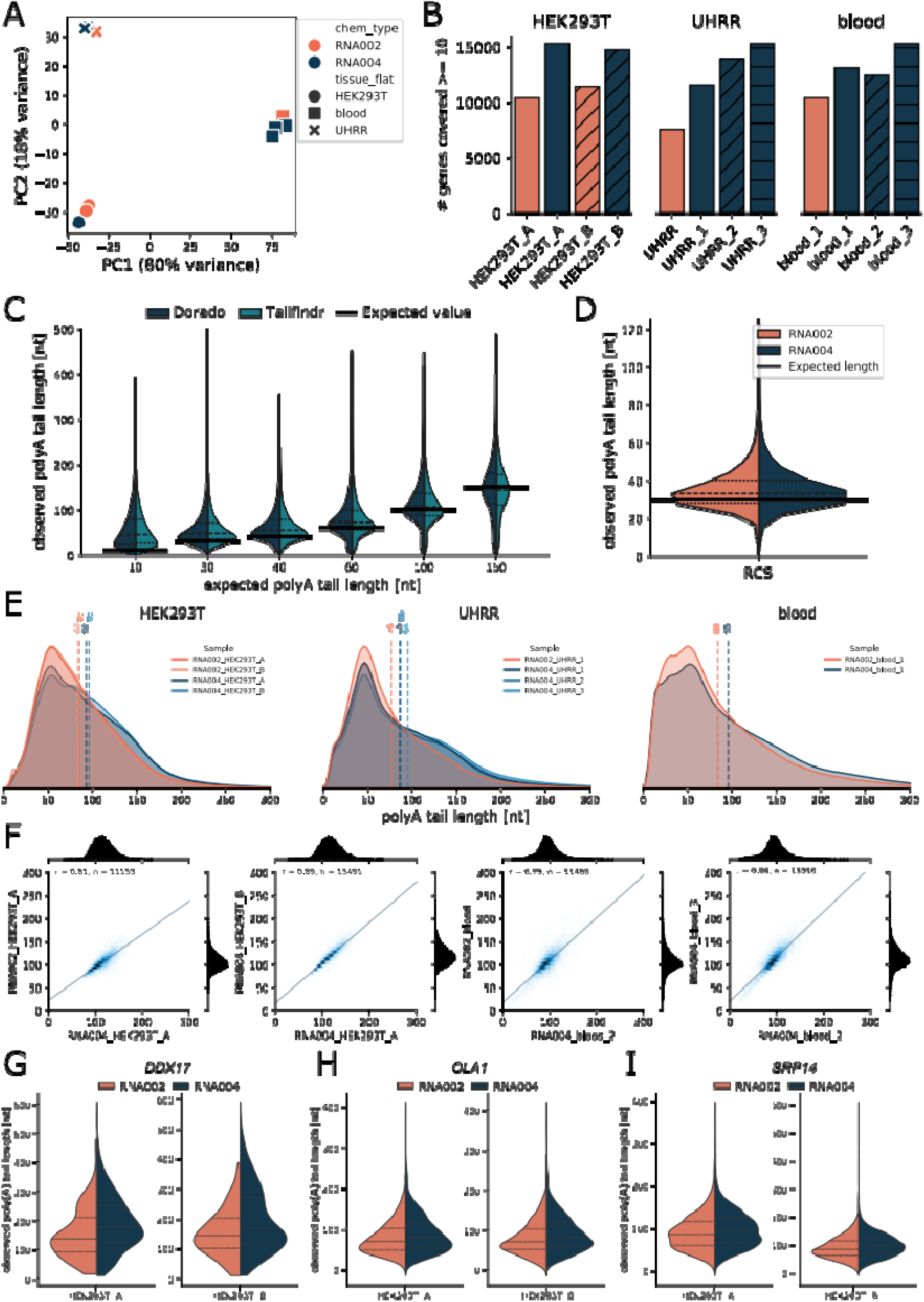
Transcriptome coverage and estimation of poly(A) length. Samples sequenced with the RNA002 chemistry and RNA004 chemistry are shown in orange and blue respectively. (A) Principal Component Analysis (PCA) based on normalized gene counts. (B) Number of genes covered with at least 10 reads (as per GRCh38 & Gencode v43). (C) Comparison of poly(A)-tail lengths from tailfindr and Dorado (0.7.2) on a RNA002-sequenced test sample. (D) Comparison of the poly(A) length distribution from the RNA Control Strand (RCS) sample with a known length of 30, sequenced with RNA002 and RNA004 chemistry (Dorado 0.7.2). (E) Mean poly(A)-tail length distribution per Gene with a minimum coverage of 10 reads. The mean values are shown as dashed lines . (F) Correlation of poly(A) lengths between genes with a minimum coverage of 10 between RNA002 and RNA004. (G–I) Poly(A) lengths comparing RNA002 and RNA004 chemistry for the genes DDX17, OLA1, and SRP14.

Furthermore, we examined the gene body coverage from the 5’ to 3’ end and observed similar coverage across all samples sequenced using either chemistry (Figure S1A).

Of particular interest for clinical applications is the coverage of genes associated with Mendelian disorders (the Mendeliome). In our analysis, the proportion of distinct disease-associated genes with at least 10X coverage ranged from 40% up to 75%, with variation depending on the sample type (Figure S1B, C, Table S2). The native human blood sample, as example of a routine diagnostic tissue, covered nearly 60% of disease-associated genes when sequenced with RNA004 chemistry.

### Evaluation of estimates of native poly(A)-tail length

Altered polyadenylation patterns on mRNA molecules are linked to various diseases, including cancer and neurological disorders, and can serve as diagnostic biomarkers or as potential therapeutic targets through their effects on gene regulation and protein expression(35, 36).

DRS can detect the length of a poly(A)-tail natively via the length and duration of the raw signal pattern for poly(A). Tailfindr was one of the first tools developed for estimating poly(A)-tail length from DRS data sequenced with RNA002 chemistry(37). Recently, ONT integrated such functionality into their production base-caller Dorado for both RNA002- and RNA004-derived data. As part of our study, we evaluated the estimation of poly(A)-tail length by ONT’s base-caller Dorado in comparison to tailfindr and examined the difference in quantity of polyadenylation between the old and new chemistry in our samples.

A RNA002 test data set with known poly(A)-tail lengths (10–150 bp) was obtained from the tailfindr publication and re-base-called with poly(A)-tail estimation using Dorado version 0.7.2. Dorado revealed similar results to both the original tailfindr assessment and the empirical label (Figure 3C).

In order to compare a standardised sequence of fixed poly(A) length between RNA002 and RNA004, we made use of a spike-in of the reference control strand (RCS), that is part of the ONT RNA kits. This sequence has a known poly(A) length of 30 nt. There was no difference in poly(A) length between RNA002 chemistry and RNA004 chemistry, both samples performed as expected (Figure 3D).

In our samples, the estimation of poly(A)-tail length by Dorado revealed similar distributions across all samples and chemistries (Figure 3E; Figure S1D-F).

Next, the correlation of poly(A) lengths was observed between RNA002 and RNA004 chemistry. The highest correlation, r of 0.89, was found between the two replicates of HEK293T cells and RNA004 chemistry, whereas the lowest r of 0.79 was between a replicate of peripheral blood in the RNA002 chemistry vs the RNA004 chemistry (Figure 3F).

Moreover, we examined the estimation of poly(A)-tail length for the genes *DDX17*, *SRP14*, and *OLA1* with long, middle, and short poly(A)-tailed transcripts, respectively. A comparison between RNA002 and RNA004 shows high concordance for all three polyadenylation patterns (Figure 3 G–I).

### Modifications detected in RNA004 samples reveal multiple uniquely modified sites

With the release of the RNA004 chemistry, the functionality for detecting m^6^A modifications in DRACH motifs and transcriptome-wide Ψ was integrated into Dorado. ONT discontinued the RNA002 kit in March 2024, which explains the absence of a production modification calling model for this version. Nevertheless, third-party tools for RNA modification detection using RNA002 have been developed and considerably advanced the field of epitranscriptomics in the past(38–42).

### Threshold definition for methylation cutoffs in RNA004 chemistry

Before the modification sets were overlapped, we defined appropriate modification thresholds. The first threshold to be defined is the modification probability on a per base level, meaning at which percentage value of modification probability is a base classified as either m^6^A or pseudouridine. A suitable starting point is to look at the histogram distributions of the modification probabilities between the native samples and the IVT samples, that are devoid of base modifications (Figure S2). As one can see from Supplementary Figure S2, the default threshold choice of the 10th-percentile would leave a high number of falsely modified reads in the IVT samples. IVT and native samples start to diverge towards the end of the distribution and consequently a threshold of 0.98 was chosen. We would like to note at this point, that this filter likely also looses some true methylation at modified sites, meaning the average percentage of site-specific methylation could be theoretically higher. But we chose this threshold with regard to the number of false positives remaining in the IVT sample.

The subsequent thresholds to be defined are the coverage threshold and the site-specific methylation threshold. To this end, we made use of our DRS samples from the peripheral blood and the orthogonal GLORI measurements that were taken from the blood of the same proband. Depicted in Supplementary Figure S3 A is the F1-score of the average methylation ratios between the GLORI samples and the DRS samples. The maximum value of the F1-score is reached at a coverage of 10 and a sitespecific modification percentage of 10.

### RNA004 vs RNA002 m^6^A predictions on chromosome 20

We first compared the performance in detecting m^6^A in RNA002 and RNA004 samples. For the RNA004 samples, we used Dorado-based m^6^A calling, whereas for those sequenced with the older RNA002 chemistry we utilized two community-developed m^6^A-detection tools: mAFiA and m6ABasecaller (see Materials and Methods, Figure 4A, Table S3 and Table S4)(43, 44).

**Figure 4.**
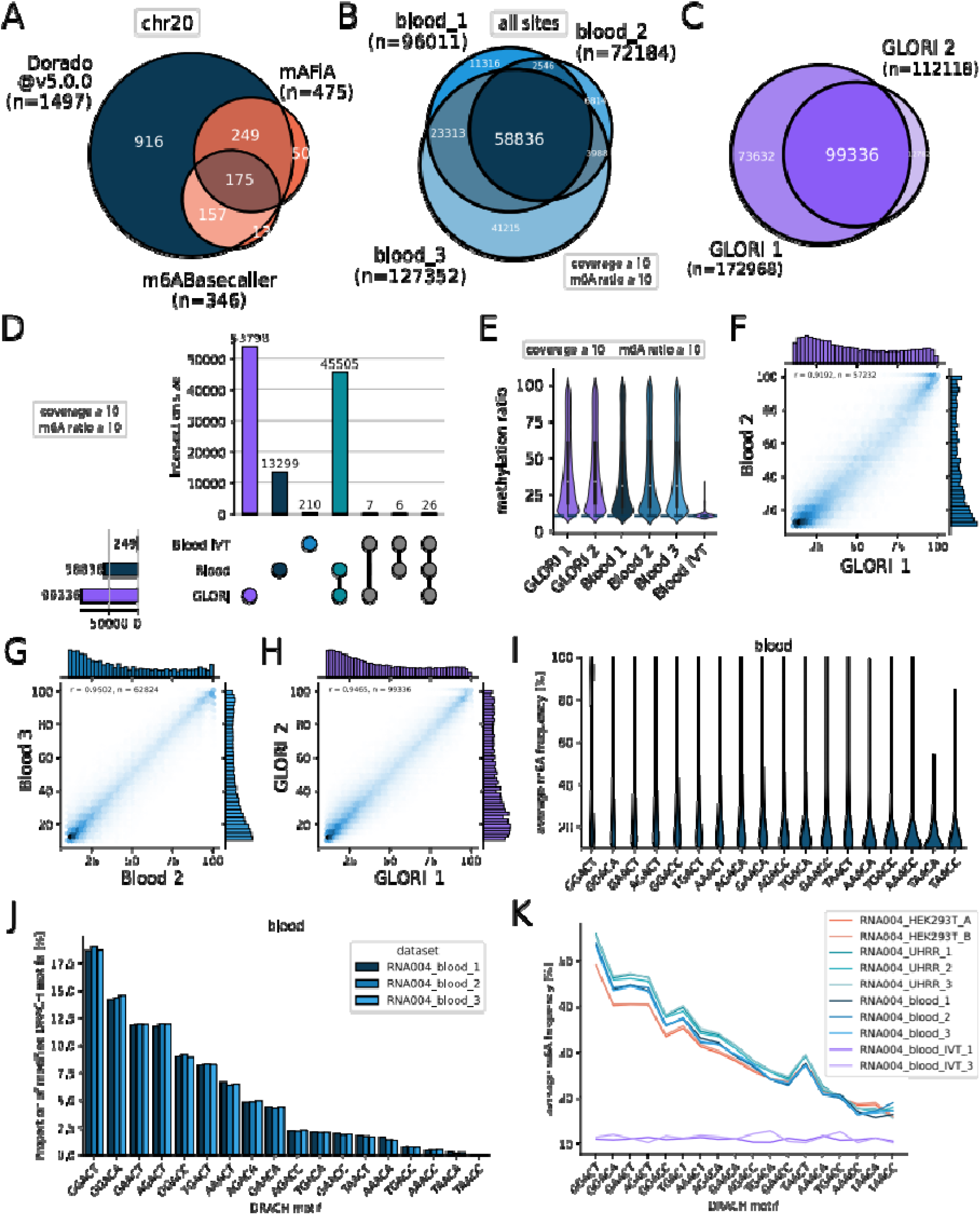
Transcriptome-wide analysis of m^6^A-sites and comparison with GLORI-sequencing as a orthogonal method of m^6^A detection. (A) Overlap between joint m^6^A-sites of blood replicates identified by RNA002 and RNA004 using mAFia and the m6ABasecaller. Only sites that are covered in RNA004 and RNA002 samples with a minimum of 10 reads are shown. (B) Overlap of joint covered m^6^A-sites for all 3 replicates of RNA004 dRNA-seq blood, which exceeded a m^6^A frequency of 10%. (C) Overlap of m^6^A-sites between two blood replicates using GLORI-sequencing. (D) Upset Plot of m^6^A-sites detected by GLORI-seq and dRNA-seq. Only sites which were covered in all dRNA-seq were kept, intersection was then performed on sites exceeding 10% modification frequency. (E) Distribution of m^6^A ratios of joint covered sites between GLORI-seq and dRNA-seq blood replicates and blood IVT without a frequency filter. (F-H) Correlation between GLORI-seq and dRNA-seq blood samples of joint covered m^6^A-sites with a minimum of 10% m^6^A frequency. The number of joint covered sites (n) and Pearsońs correlation coefficient (r) are shown in the plots. (I) Mean m^6^A frequency per site of all blood samples per DRACH motif. (J) Proportion of modified DRACH motifs compared to all modified DRACH motifs of three blood replicates. (K) Average m^6^A frequencies per DRACH motif and sample.

Using RNA004 samples in combination with the ONT base caller detected the highest number of m^6^A sites (1497), whereas applying m6ABasecaller and mAFiA to RNA002 samples resulted in significantly fewer m^6^A sites, namely (346) and (475); measurements were made for chromosome 20 (Figure 4A). Because the new chemistry translates sample strands faster through the nanopore, we were also interested in modified sites detection, when correcting the RNA004 samples for coverage. The individual blood replicates RNA004 still detected more m^6^A sites than the legacy basecallers (Table S4).

### Detection of modifications in a blood sample as an exemplary clinical tissue

Next, we investigated the performance of m^6^A and Ψ base calling by Dorado in the 3 replicates of RNA004-sequenced native blood sample of a healthy individual compared to its two corresponding unmodified IVT samples and the two GLORI replicates for orthogonal m6A verification. In the filtered native blood samples 96.011, 72.184 and 127.352 m^6^A sites were detected. For the unmodified IVT sample, the number of probably false-positive predicted m^6^A sites was 249, which is, however, only a minor fraction of the natively modified counterpart. The GLORI samples had 172.968 and 112.118 m6A sites (Figure 4B, C, Figure S3 B and Table S4).

The intersection of all 3 sample types has the highest overlap between blood and GLORI, with 13.299 sites in just blood DRS and 53.798 sites in just GLORI (Figure 4D).

When examining the methylation ratio across all sites, we observe a bi-modal distribution, with the IVT having a lower average methylation than the native samples (Figure 4 E).

The cross-correlation of sites shared between replicates of native blood or GLORI can be seen in Figure 4 F-H and Supplementary Figure 4 C-H. And we observe that replicates of the same technique for example, GLORI 1 vs GLORI 2 show a higher correlation than replicates of GLORI vs DRS blood.

By examining the average m^6^A frequency across modified DRACH motifs, we observed that the natively modified blood, UHRR and HEK293T samples have a characteristic distribution as reported in literature(32, 39, 45). In contrast, the frequency of false-positive detected m^6^A sites in the unmodified IVT sample was evenly distributed across the DRACH motifs (Figure 4I-K; Figure S6 and Table S5).

Furthermore, we were interested in the number of modification sites in genes associated to Mendelian disorders, the so-called “Mendeliome”, which can be predicted by the Dorado m^6^A caller. From all m^6^A sites detected in either replicate of a healthy blood sample 30585, 23627 and 39257 were in Mendeliome genes, respectively (Table S3 and Figure S3 K). When requiring the intersection of 3 of 3 replicates and the Mendeliome gene set, 19308 sites were found. Potential aberrations of modification stoichiometry in these regions might influence the function of gene products, their physiology and pathophysiology.

The transcriptome-wide Ψ scan by Dorado in the three replicates of the native blood sample predicted the existence of 30163, 23456 and 47435 potential modification sites with a modification frequency of at least 10% and a valid coverage of 10 reads. For comparision the two IVT replicates had 10477 and 14420 sites each (Fig 5A, Table S6 and Table S7). In the intersection between 3 of 3 replicates of peripheral blood 4555 sites remained, most sites were unique to the respective biological replicates (Fig 5B). Supplementary Figure 5A shows the remaining overlaps after asking for joined sites and Figure 5B after subtracting the IVT.

**Figure 5.**
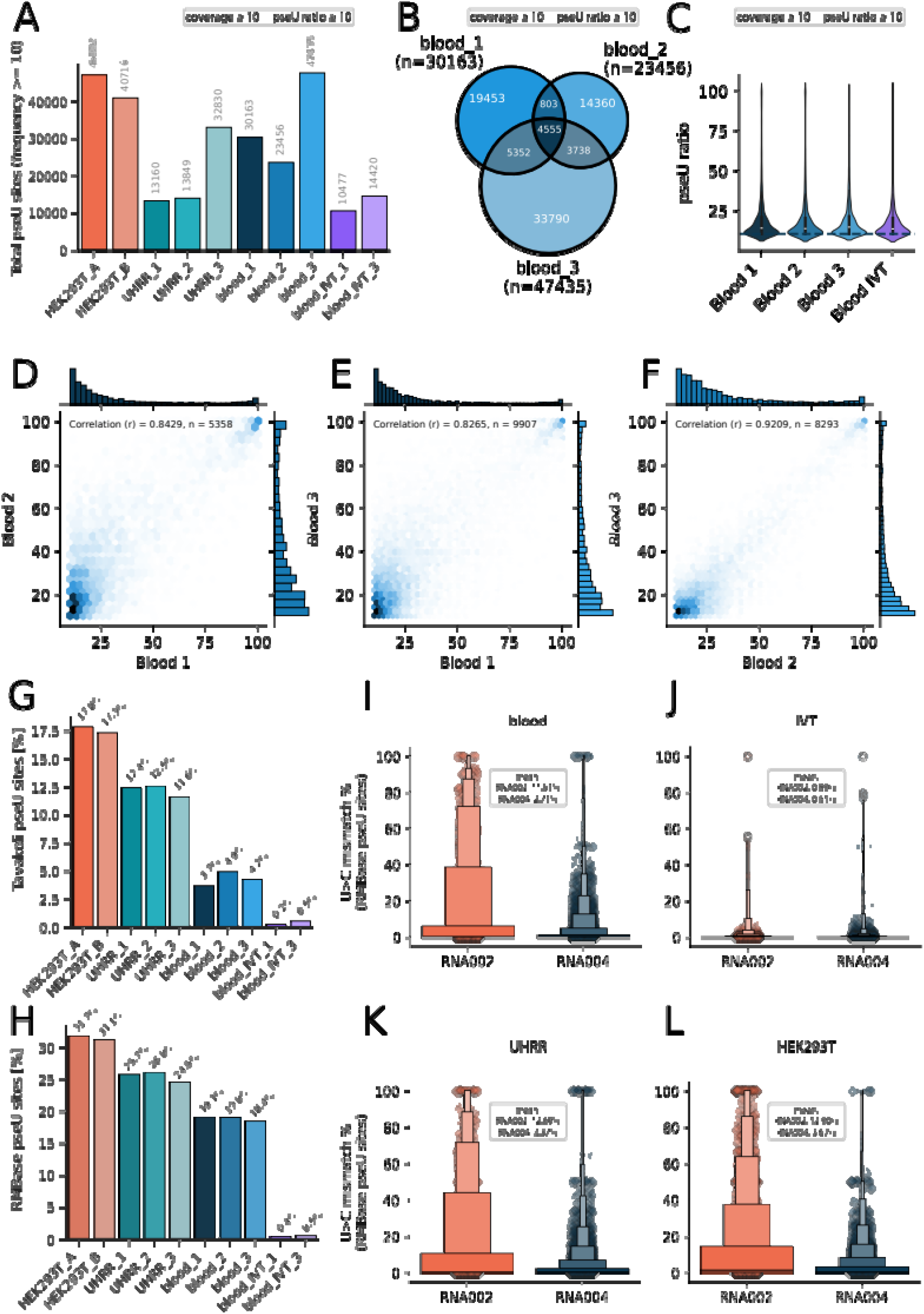
Transcriptome-wide analysis of Ψ sites detected by DRS. (A) Number of detected Ψ sites per sample with a minimum coverage of 10 and minimum frequency of 10. (B) Overlap of all detected Ψ sites between DRS blood replicates which exceeded a Ψ ratio of 10 % with a minimum coverage of 10. (C) Global distribution of Ψ ratios between replicates of all detected sites by DRS with a minimum of 10 percent Ψ frequency (dashed line). (D-F) Correlation of joined covered Ψ sites between blood replicates with a minimum of 10% Ψ frequency.The number of joined covered sites (n) and Pearsońs correlation coefficient (r) are shown. (G) Proportion of detected Ψ sites compared to all listed Ψ sites provided by Tavakoli et al.(46).(H) Proportion of detected Ψ sites compared to all listed Ψ sites provided by the database RMBase[(47).(I-L) Distribution of U>C mismatch frequencies of Ψ sites provided by RMBase in different samples with the respective mean.

The methylation ratio between the three native DRS samples and the IVT is highly similar and most sites are close to the methylation filter cutoff rate of 10 % (Fig 5C; Fig S 5C). This in contrast to the m^6^A sites, where the average methylation was a bit higher and in the m^6^A samples there was a marked contrast between the three native samples, with the average methylation ratio bein much lower in the IVT sample. Likewise, the cross-correlation between the biological replicates ranges from r = 0.82 to r = 0.92 and is a bit lower than for the m^6^A sites as well (Fig 5 D-F). This is consistent with the fact that low stochiometry sites are less likely to replicate and more noisy.

Approximately 1633 pseudouridylation sites were found to be covered in 3 of 3 replicates in the Mendeliome gene set (Fig S 5 D, E). Of the > 60, 000 stop-codon sites in the human transcriptome, ∼1% were predicted to be modified in at least one replicate. However, only 10 sites remained when asking for the intersection of all 3 replicates (Fig S5 F, G). Pseudouridylation can trigger protein readthrough, making these predicted modification sites an interesting target for investigating premature termination codons (PTCs)(4).

Given the recent reports precedence for U>C mismatch at Ψ sites, we wondered how this would manifest in a transcriptome-wide manner(46). First, we determined the ratio of detected Ψ sites vs all sites in the set of high-confidence sites published by Tavakoli and co-workers and sites detected in RMBase for all samples (Figure 5 G-J, Table S6, Table S8, Table S9)(47). In the native blood samples, around ∼4-5% Ψ sites were detected. For the IVT only 0.2-0.5% of sites were found. Just for comparision, around 17% of Ψ sites are found in the HEK293T samples and around 12 % of Tavakoli sites are found in the UHRR samples (Fig 5 G). When looking at RMBase sites, blood had around 20% of Ψ sites detected. The HEK293T samples had > 30% of RMBase sites detected (Fig 5H). Figure 5 I-J show the U>C mismatch percentage for covered RMBase sites in the three native samples and the IVT. Expectedly, the older RNA002 chemistry had higher mismatch percentages throughout all sample classes, ranging from ∼ 13% in the HEK293T samples to ∼ 1% in the IVT. In contrast, the newer RNA004 chemistry only had 3.67 % error rate in HEK293T samples and only 0.51 % error rate in the IVT. Thus, showcasing the overall lower error rate for RNA004 in the context of U>C mismatches and indicating diminishing returns for misbasecall counting based strategies in the face of improved kits and basecaller models.

### Transcriptome-wide cross-correlation of GLORI m6A with DRS HEK293T data

Next, GLORI sequencing was repeated in-house for three replicates of HEK293T cells (see Table S4). Overall, RNA004 chemistry on the P24 in connection with the new Dorado model recovered the most m6A sites, followed by the in-house GLORI data. Overall, the cross-correlation was naturally higher in replicates of the same type and ranged from R²=0.9571 for the to GLORI replicates to R²=0.8862 for an overlap between HEK293T DRS and GLORI.

### RNA004 accurately reads the pseudouridylation stoichiometry in a targeted reporter system

Next, we investigated detecting modifications in a site-specific manner, since another question is whether the technique is mature enough to track and target known positions in order to be developed into a clinical assay.

First, validated the Ψ stoichiometry determined from a custom targeting pseudouridylation system developed by Schartel and co-workers(6). Three HEK293T samples were transfected with a modified pseudouridine synthase (DKC1), artificial guide snoRNAs as well as a selectivity reporter sequence containing EGFP and mCherry sequences, each harboring a target motif that represents a premature stop codon. In sample A, both EGFP and mCherry motifs are expected to be equally targeted for pseudouridylation, whereas in sample B, the mCherry motif is preferentially targeted and therefore expected to be modified to a greater extent than EGFP (Figure 6A and Figure S7). Sample C contained a scrambled guide RNA with no pseudouridylation capability and is used as an unmodified biological control.

**Figure 6.**
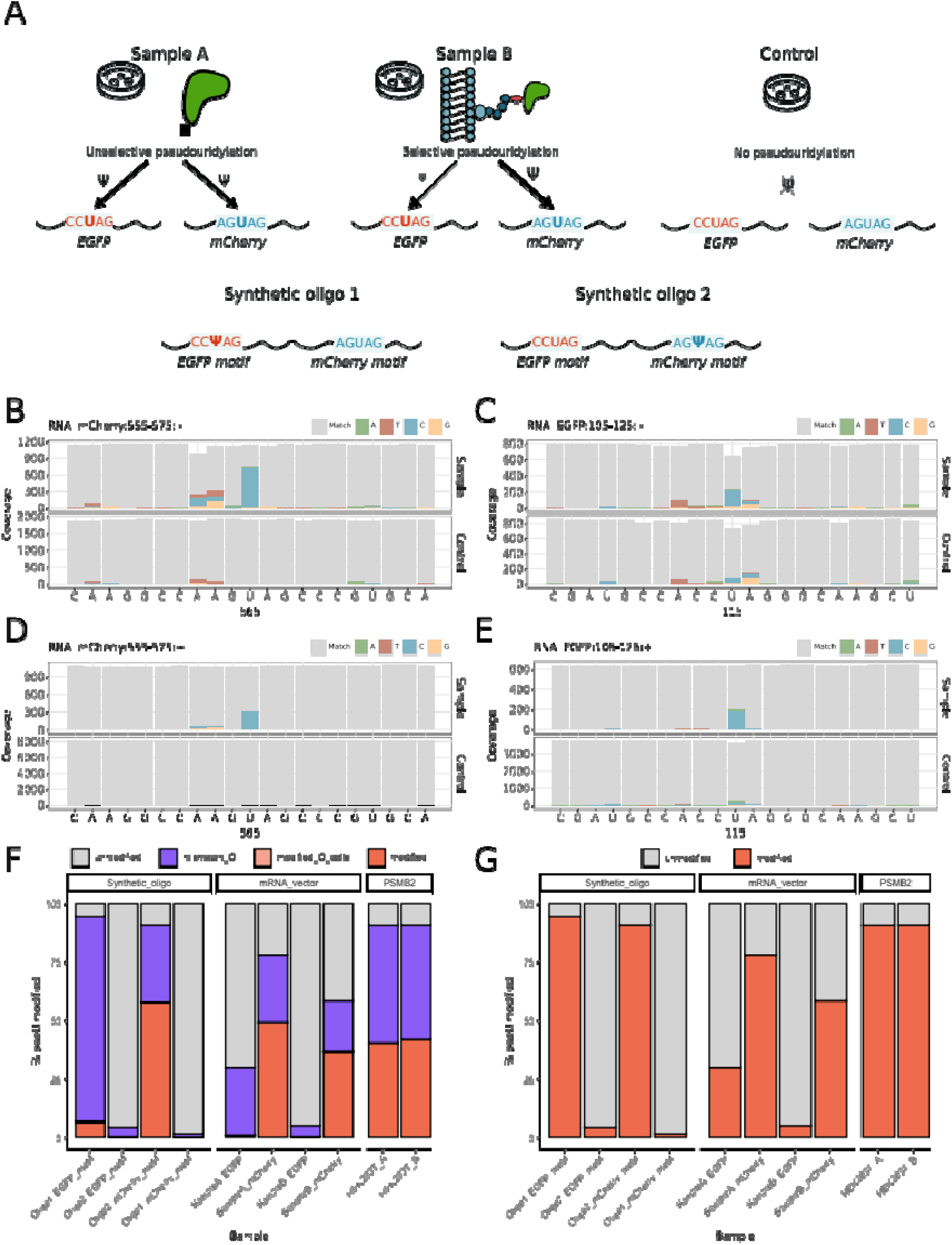
Site-specific Ψ modification. (A) Overview of the sample types. Condition A: EGFP and mCherry were both targeted equally for pseudouridylation after transfection into HEK293T cells; condition B: mCherry was preferentially targeted; control: neither mCherry nor EGFP were targeted, and in addition two oligos were sequenced with 100% motif-specific Ψ as a modification control. (B,C) C–U mismatch at Ψ sites using mCherry or EGFP with RNA002 chemistry and HEK293T (Condition A). (D,E) C–U mismatch using RNA004 chemistry and condition A. (F) Percentage barplot of U classified as Ψ, including the canonical U miscalled as C (green) and miscalled C that the Dorado model recognized as Ψ (blue). (G) Percentage barplot of U classified as Ψ (red) and classified as canonical U (gray) for vector samples, oligos, and HEK293T cells, after merging both miscalls and mod predictions.

First, we checked whether the targeted sites in the mCherry and EGFP mRNAs can be detected by both RNA002 and RNA004 chemistry by comparing the base-calling errors between targeted samples A and B to those of the non-pseudouridylated sample C using nanoCEM(48)

Next, we evaluated the performance of the Dorado-based Ψ detection for RNA004 based on 1) synthetic oligos that contain the EGFP and mCherry motifs of the targeted reporters both in fully modified and unmodified states; 2) the pseudouridylation-targeted sites of the EGFP and mCherry reporters in sample A and B; and 3) a high-confidence Ψ site in the *PSMB2* transcript of HEK293T cells (Figure 6A, see Materials and Methods)(49).

For both unmodified motifs in the synthetic oligos, the number of false-positive detected Ψ was rather low, with less than 2.5% of reads modified as determined by Dorado (Figure 6F, Table S10). The fully pseudouridylated mCherry motif revealed 60.69% modified reads, whereas for the EGFP site with the same expected Ψ stoichiometry, only 10.20% of reads were found to be modified (Figure 6F). Interestingly, the EGFP motif reveals a particularly high U>C mismatch rate (∼90% of reads) and as Dorado detects modifications at read level, considering only U-sites, C-called bases are neglected. When combining the number of C-mismatches with the number of modified reads called by Dorado both on U and C bases (Figure 6 B–G, Table S10; see Materials and Methods), the percentage of modified reads increases to 98.38% and 93.77% for the positive controls of the EGFP and mCherry motifs, respectively (Figure 6G).

The same pattern was observed for the targeted EGFP and mCherry reporters in samples A and B and only if both the C-mismatch and base-caller-derived Ψ sites were used can the expected differences in stoichiometry between samples A and B be verified(6). Specifically, the modification frequency ratio between mCherry and EGFP was higher in sample B (9.6-fold) compared to sample A (2.6-fold).

Moreover, the high-confidence Ψ site in *PSMB2* transcripts reveals similar modification frequencies in both HEK293T samples, and by using both C-mismatches and Dorado-called Ψ sites, the modification frequencies were 11% above the expected stoichiometry of 80% modified reads reported in literature (49).However, the number of Ψ sites discovered by Dorado amounted to only 40 and 42% of reads, respectively, which can be explained by Dorado’s basecalling and modification model architecture.

### Observations in Reference Oligos used for ONT model training

Recently, ONT has made a set of oligos available, that were used for internal training and validation (Figure S8 and S9). In order to show the impact of the selected modification probability cutoff in this paper, Figure S8 A shows the unmodified control strand without any filter preset. That means any modification probability from 0-1 will be shaded. After applying the chosen filter threshold of 0.98 modification probability, as derived from the histogram data, the same control oligo is now free of methylated bases (Figure S8 B). We acknowledge, that this filter cutoff might loose true methylation at some sites. If orthogonal mass-spectroscopy data at a known site is available, one may even relax the filter threshold in order to keep higher stochiometries. In Panel S8 B, we observe a false positive pseudouridine call in the m6A reference data. This false positive site will naturally remain unaccounted for, if just IVT is subtracted from the modified sites and illustrates a problem of some current approaches for estimations of total accuracy. In Figure S9 A, we observe around 60% C/U mismatch at a known pseudouridine site. For our data, in the fully modified EGFP motif of the oligo 1, Dorado predicts an m^6^A modification directly next to the Ψ site (+1), which should not be present in the synthesized sequences (Figure S9 B). This shows that the performance of Dorado-based modification detection is dependent on the sequence context. And generally we would urge caution that the accuracy values defined on these oligos are true for the respective test sets, but will not neccessarily transfer into a biological system.

### Putative loss of function in METTL5 and site-specific m^6^A detection

Finally, we present a clinical case from the Institute of Human Genetics Mainz for which we were able to validate the functional impact of genetic variations within a methyltransferase gene that is responsible for a site-specific m^6^A modification using DRS.

A one-year-old girl showed severe microcephaly (occipito-frontal head circumference > −6 standarddeviation) and developmental delay. Two compound heterozygous variants, c.224+5G>A (p.(?)) and c.427A>T (p.(Lys143*)), in the *METTL5* gene (NM_014168.4) were identified by whole exome sequencing and suspected as the underlying cause for an autosomal recessive intellectual developmental disorder type 72 (OMIM # 618665) (Figure 7A–C).

**Figure 7.**
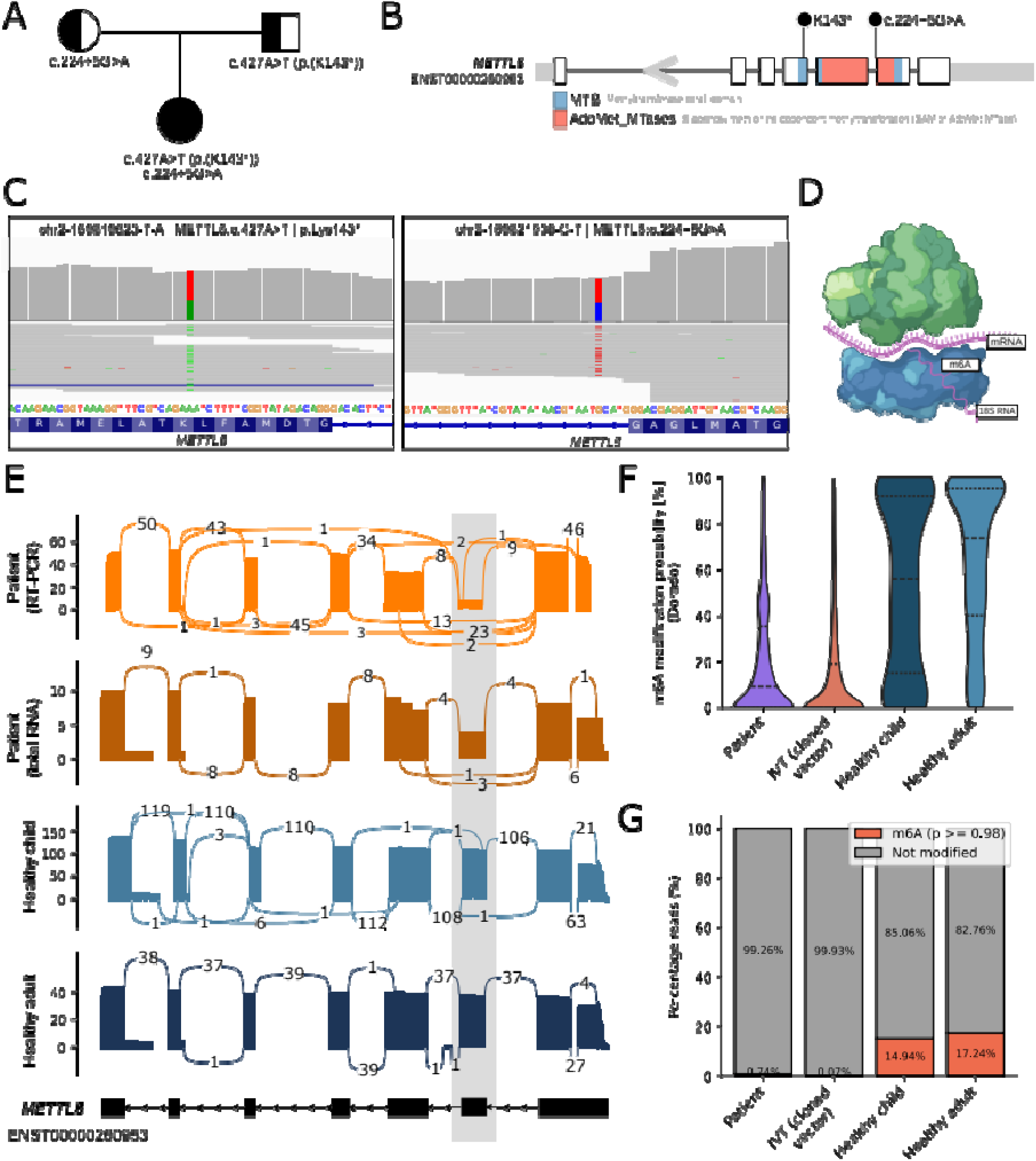
Site-specific m6A and mutations detected in METTL5 in the patient by DNA sequencing. (A) Family tree of the heterozygous parents and the compound heterozygous child. (B) METTL5 gene overview with location of specific mutations. (C) Mutations on the DNA level (IGV). (D) Overview of the ribosome and m6A at position 1832 close to the active site. (E) METTL5 splice assays for controls and patient, the mis-spliced exon is highlighted by the shaded box. (F) Violin plot of modification probability, as estimated by the base-caller, at position A1832. (G) Percentage of reads which were either identified as unmodified or modified (with a probability threshold of 0.98) at position A1832 of the 18S rRNA in the patient, healthy control samples and the IVT of a cloned vector.

While the nonsense variant c.427A>T (p.(Lys143*)) in exon 4 of the *METTL5* gene was classified as pathogenic based on ACMG guidelines (pathogenicity level 5, PVS1+PM2=8+2=10 points), the intronic variant c.224+5G>A was classified as a variant of unclear significance (VUS, pathogenicity level 3, PP3+PM2+PM3=1+2+2=5 points). To investigate the effect of the VUS on *METTL5* splicing, the RNA extracted from peripheral blood of the patient was sequenced using DRS (RNA004) and revealed skipping of exon 2 in approximately 50% of the reads, suggesting a loss-of-function splicing defect. To validate this analysis and increase the vertical coverage of *METTL5* transcripts, a targeted RT-PCR assay of exons 1–7 of the *METTL5* transcript was applied and additionally confirmed the aberrant splicing pattern (Figure 7A and Figure 7E).

Since *METTL5* is known to elicit an m^6^A modification not globally but at a single site that is close to the active site of the ribosome (Figure 7D), we were interested whether we could verify aberrant m^6^A modification in the peripheral blood of this patient. The patient sample showed reduced m^6^A modification at the METTL5 target position 1832 of the 18S rRNA compared to healthy pediatric and adult samples (Figure 7F, G). For the first time, we can confirm the loss-of-function of an RNA modifying enzyme in a clinical case via DRS.

## Discussion

DRS has revolutionized RNA analysis by enabling the detection of both full-length transcripts and transcriptome-wide modifications from native RNA molecules. This approach has the potential to deepen our understanding of the complex epitranscriptome, which encompasses over 170 known modifications. The recently released RNA004 chemistry, featuring new base-calling models and integrated capabilities to detect modifications such as m^6^A and Ψ offers exciting prospects for establishing DRS as a routine tool in both epitranscriptomic research and clinical applications.

### RNA 002 versus RNA004

In this study, we assessed the RNA004 chemistry compared to the earlier RNA002 version, focusing on improvements in sequencing quality, throughput, and novel capabilities for detecting modifications and demonstrated the first clinical application of DRS.

In 2019, the first study to comprehensively analyze DRS for a human poly(A)-selected RNA derived from a cell line utilized 30 MinION flow cells to generate 9.9 million aligned reads with a median identity of 86% and a maximum read length of 21, 000 bases(50).This effort was shared between six different institutions.

In our study, both single PromethION flow cells loaded with poly(A)-selected RNA from HEK293T cells provided each more output with higher quality at a fraction of the previous cost and effort.

However, some limitations of the older chemistry, such as read lengths and transcriptome assessment, persist (26). In particular, capturing full-length transcripts remains problematic, largely due to a mismatch between annotated transcript lengths and the fraction of reads covering the annotated 5’ end. This issue arises from the DRS adapter design and is compounded by the fact that the motor protein, which translocates RNA through the pore, eventually releases the 5’ end when the molecule is fully processed. Previous studies have shown that this results in the last few nucleotides being unsequenced, a challenge shared with the new chemistry as the adapter attaches only to one end of the RNA molecule (50). This limitation also impacts state-of-the-art transcript-detection tools, such as bambu, which struggle to accurately predict the 5’ end, even in “full-length” direct RNA reads(51). Nevertheless, special strategies for preparing sequencing adapter libraries, combined with specific changes to MinKNOW’s read detection algorithms, can partially mitigate the issue of incomplete or missing transcripts(52, 53). For a comprehensive and up-to-date overview of isoform detection, we recommend the LRGASP study(54).

### Site-specific modification stoichiometry

The integration of models for detecting m^6^A and Ψ modifications in Dorado has created opportunities to utilize DRS in routine analyses. By taking advantage of the new RNA004 chemistry and enhanced capabilities for detecting RNA modifications, we demonstrate precise estimation of site-specific Ψ stoichiometry within a targeted system used for drug development. This level of detection, achieved at single-nucleotide resolution, challenges the conventional methods of detecting Ψ. Thus, DRS is a valuable tool for the rapid and straightforward quality assessment of therapeutic RNAs, such as mRNA vaccines and antisense oligonucleotides(55). Additionally, DRS offers a comprehensive single workflow for evaluating sequence identity, integrity, poly(A)-tail length, and contamination from oligonucleotides, thereby streamlining quality control processes for therapeutic RNAs(56).

Furthermore, RNA modifications play an important role in the development of mRNA vaccines. Ψ can suppress recognition by toll-like receptors in the innate immune system. This reduces the immunogenicity of the RNA, which was a crucial breakthrough for developing effective mRNA vaccines against the SARS-Cov-2 virus with reduced side effects and improved protein translation(57, 58).

### RNA modification detection in diagnostics

We were also able to confirm the pathogenicity of variants in an RNA-modifying enzyme (METTL5) by predicting the m^6^A stoichiometry on the 18S rRNA in a clinical patient with an intellectual developmental disorder. Besides the known disease-association of aberrant RNA-modifying functionality, there is growing evidence that dysregulation of mRNA modifications contributes to tumor development and progression, making them promising targets for future drug development(59–62). Beyond mRNA and rRNA, DRS shows growing potential for studying modifications in other RNA species, including mitochondrial RNA, tRNA, and other non-coding RNAs(16, 25, 63, 64). These RNA types offer unique opportunities for elucidating disease mechanisms related to RNA-modification disorders(7). Accurate prediction of molecular changes is essential for understanding disease mechanisms, and improvements in DRS accuracy with RNA004 chemistry are poised to enhance biomedical research further.

Comprehensive analysis of the epitranscriptome will be pivotal not only for studying rare diseases but also for advancing cancer diagnostics and RNA therapies. The integration of RNA epi-signature analysis into the clinical routine screenings holds the potential to improve diagnostic precision and deepen our understanding of pathomechanisms in rare diseases. Thus, routine use of DRS in clinical settings is increasingly realistic.

### Limitations of current RNA modification tools

Although methodologies for detecting certain RNA modification are well established, current methodologies still face significant limitations. Despite the availability of several community-developed tools optimized for RNA002 chemistry, no existing method can comprehensively detect more than a few RNA modifications(41, 44, 65).L Such distinctions, however, are crucial for advancing our understanding of the regulatory functions of RNA modifications and their implications for health and disease. Reliable, user-friendly solutions to detecting RNA modification and their integration within standard software are especially needed for clinical applications and would facilitate better assessment and diagnosis of modopathies.

### Model complexity and benchmarking challenges

The development of new classification models for detecting RNA modification is ongoing. Typically, new models are benchmarked against older tools using different training data or chemistry, leading to significant model complexity and heterogeneity. This complexity can be overwhelming for practical use, as discussed in the one of the most recent and comprehensive reviews of m^6^A base-calling models to date (66).The review evaluated 14 m^6^A-detection tools but found no universal model suitable for all applications. For example, a model trained on human cell line data performed poorly on oligonucleotide data, and *vice versa*. In additon to the mere basecaller error, the accuracy of the modification in question will also be directly dependent on the ability to define exact methylation filtering cutoffs, based on external criteria. As an example, the m^6^A modification filter adjustment in this paper had several benefits in comparision to the Ψ filter adjustment, such as clearly defined DRACH-motifs vs all A and orthogonal validation via GLORI. These additional filters increase the accuracy of m^6^A detection irrespective of the basecaller model and serve as an additional explanation why Ψ detection remains challenging. This finding is also echoed by recent modification detection studies(67).

### Persistent error patterns in RNA004 chemistry

Liu-Wei and colleagues investigated systematic base-calling errors in DRS and found in canonical nucleotides that, despite improvements in accuracy, the RNA004 chemistry still exhibits similar error patterns as its predecessor, such as frequent insertion and deletion errors(29). The misbasecalls on modified nucleotides, observed in the TRID system, might arise due to the *k*-mer data used for RNA004-based training not being fully representative of all sequences. Consequently, certain sites can show base-calling errors exceeding 50%.

### Lack of gold standard data sets

In our study, we have predicted several thousands of Ψ sites using RNA004 and the Dorado basecaller, while previous publications reported only several hundred Ψ sites on mRNA level(68). However, the overlap of the four different studies was only marginal with sensitivity and specificity being unassessed. Even minor variations in these parameters can produce disparate results when calculating overlaps (68). Benchmarking the detection of RNA modification is further complicated by the absence of universally accepted gold-standard data sets, inconsistent sequencing depths, and diverse post-base-call filtering options.

Another illustrative example is the comparison of called m^6^A sites observed by the community-based m^6^A detection tool CHEUI (CH3 (methylation) Estimation Using Ionic current) using HEK293T cell line data with the GLORI dataset, a gold standard for human m^6^A sites(69). Recently, Chan and colleagues reported a site-level stoichiometry correlation with GLORI of 0.64, while the CHEUI developer itself found a correlation of 0.85 (39, 69). The HEK293T DRS vs HEK293T GLORI correlation in our study was closer to the second example.

The intricacy of the RNA epitranscriptome further complicates the generation of ground-truth data sets. With over 170 known RNA modifications, it is uncertain whether each modification leaves detectable deviations in retention time or current levels. Additionally, nanopore devices detect *k*-mers, so signals are often influenced by adjacent bases within the sensing zone, potentially leading to false positives. This might explain erroneous m^6^A detections at the +1 position of Ψ in oligonucleotide sequences in this study. Moreover, several modifications in proximity are difficult to resolve and may require specific enzyme knockouts, which further increases data set complexity. For instance, 19 modifications in *E. coli* tRNAs separated by fewer than five nucleotides required methyltransferase knockdowns to isolate their signatures(70). And we would also raise caution when transfering accuracy values derived from a control data set with a limited number of oligos into a biological system that the biological distribution and relevance must be taken into account. That means, if a highly present modification, such as m6A, causes even 1-2% false positives in a control data set and the false positives are a very lowly abundant modification. Then, the model error for the lowly abundant modification may be as high as the number of true sites. Even if the pre-computed model error seems small in the control data set. All in all, this makes generating ground-truth data sets and benchmarking challenging.

### Toward broader clinical application of direct RNA sequencing

For DRS to achieve widespread use in detecting RNA modifications in clinical settings, the development of gold-standard data sets for human samples, such as those established by the Genome in a Bottle (GIAB) or the Challenging Medically-Relevant Genes Benchmark-Set (CMRG), is essential(71, 72). Another current limitation is the absence of ONT-based barcoding kits for RNA004 chemistry. This forces users to sequence an entire flow cell per sample or resort to a “nuclease flush” to remove libraries from the flow cell.

### Conclusion

Despite these challenges, RNA004 chemistry offers significant improvements in sequencing accuracy and throughput. Site-specific detection of modifications holds promise for integration into clinical practice, with applications extending beyond m^6^A and Ψ to other modifications. Potential uses include sitespecific assays and quality control of RNA therapeutics, as we could demonstrate in our paper given the TRID system and the *METTL5* case. The growing number of RNA004 preprints is already encouraging and signifies heightened interest in the new kit(68, 73). A larger userbase could provide the impetus to close these gaps and ultimately realize the potential of DRS to enrich clinical care and diagnostics.

#### Ethical statement

This is a basic research project to validate direct RNA sequencing to evaluate its suitability for detecting molecular targets for innovative forms of therapy. In addition to a standardized sample (blood from a healthy volunteer), we also examined blood from an infant with an autosomal recessive intellectual developmental disorder type 72 showing a putative splice site on *METTL5* and the observed reduction of m^6^A at the A1832 position of 18S rRNA in the patient. Since the infant was unable to understand the aims, scope, risks, and benefits of the study, and because we are reporting on a rare disorder, the patient was considered highly vulnerable. Thus, informing the parents as legal proxies about all aspects of the rather complex procedure was paramount to safeguard the interests of the patient. The diagnosis and the associated functional consequence were determined in one assay using nanopore sequencing. Regarding its suitability for the detection of molecular markers, artificial modifications of RNA were also used to test the stability, sensitivity, and selectivity of the method for identifying pathologically relevant molecular targets.

The project was evaluated by the internal ethics advisory board of the University Medical Centre. From an ethical point of view, this is basic research without direct reference to patient care. Informed consent was obtained from the legal proxies that surplus material (blood) was intended to be used for the validation of a new method for direct RNA sequencing. Data was anonymized and the risk of reference back to individuals due to the processing of genetic information in the case of rare disease was pointed out, as well as the fact that no whole genome data was generated or analyzed. However, both the proband and legal proxies consider the possible future risk to be acceptable when weighed against the gain in knowledge.

The research presented here is explicitly not a clinical study. The study was therefore evaluated by the internal ethics advisory board. Ethical principles, in particular the principle of autonomy, are upheld, which is especially true in light of the revision of the Declaration of Helsinki, which aims to enable research in this area while maintaining the protection of vulnerable groups such as children in order to facilitate access to innovative medical procedures. This also applies in the case of the present study to validate the clinical applicability of new diagnostic procedures or the identification of molecular targets, even if there is currently no direct patient benefit but at most a group benefit. This study, using a single sample of one vulnerable patient providing relevant information diligently and obtaining fully informed consent of the legal proxies to validate a novel diagnostic strategy does not raise ethical concerns. However, should the concept be translated into a (translational) clinical study, ethical approval would have to be obtained by the regulatory authorities.

For the healthy control the use of blood data received ethical approval, granted on April 9, 2025, by the Landesärztekammer Rheinland-Pfalz (Reference number 2025-18081). Informed consent was obtained from all participants.

## Data Availability

The data for this study have been deposited in the European Nucleotide Archive (ENA) at EMBL-EBI under the accession number PRJEB74238. The human phenotype data will be deposited to EGA once the manuscript has been conditionally accepted.

## Code Availability

All code written in support of this publication is publicly available at https://github.com/CSG-Group-Mainz/RNA004-Manuscript.

## Acknowledgements

This work was partly funded by Deutsche Forschungsgemeinschaft (DFG, German Research Foundation; project no. 439669440 TRR319 RMaP TP A01/A05/C01/C03 to F. L., J. K., M.H. and S.M). S.W. and S.G. acknowledge funding from the Emergent AI Center funded by the Carl-Zeiss-Stiftung. S.S. and S.G. acknowledge funding from the Forschungsinitiative Rheinland-Pfalz and the ReALity initiative of the Johannes Gutenberg University Mainz. S.Sy. acknowledges the M3odel initiative from the Forschungsinitiative Rheinland-Pfalz. This work was also partly supported by funding from ERC ADG MultiOrganelleDesign (E.A.L.). S.G. and C.H. acknowledge funding from the Boehringer Ingelheim Stiftung.

F. L. and J. K. thank the Next Generation Sequencing Core Facility of the German Cancer Research Center, particularly Franziska Petermann and Panagiotis Provataris for their support.

## Author Contributions

C.H. designed the project, wrote the manuscript, and performed data analysis. A.W. performed data analysis, wrote the manuscript, and composed the figures. S.D., V.H., F.K., and L.H. supported with patient recruitment and clinical interpretation of the variants. T.B. performed the sequencing of the cell line data. J.F. sequenced the peripheral blood samples supported by K.J. S.M. sequenced the oligos under the supervision of M.H. J.M., S.S., V.D., K.B., S.W., and F.H. contributed to data analysis, to writing the manuscript and designed parts of the figures. L.S. designed the TRID system and performed the analysis under the supervision of E.A.L. J.K. prepared the GLORI sequencing data under the supervision of F. L. S.G., and M.L. supervised the study, edited the manuscript and contributed to writing and conceptualizing the manuscript. All authors approved and proofread the manuscript.

