## Supplemental Information for "Direct RNA sequencing enables improved transcriptome assessment and tracking of RNA modifications for medical applications"

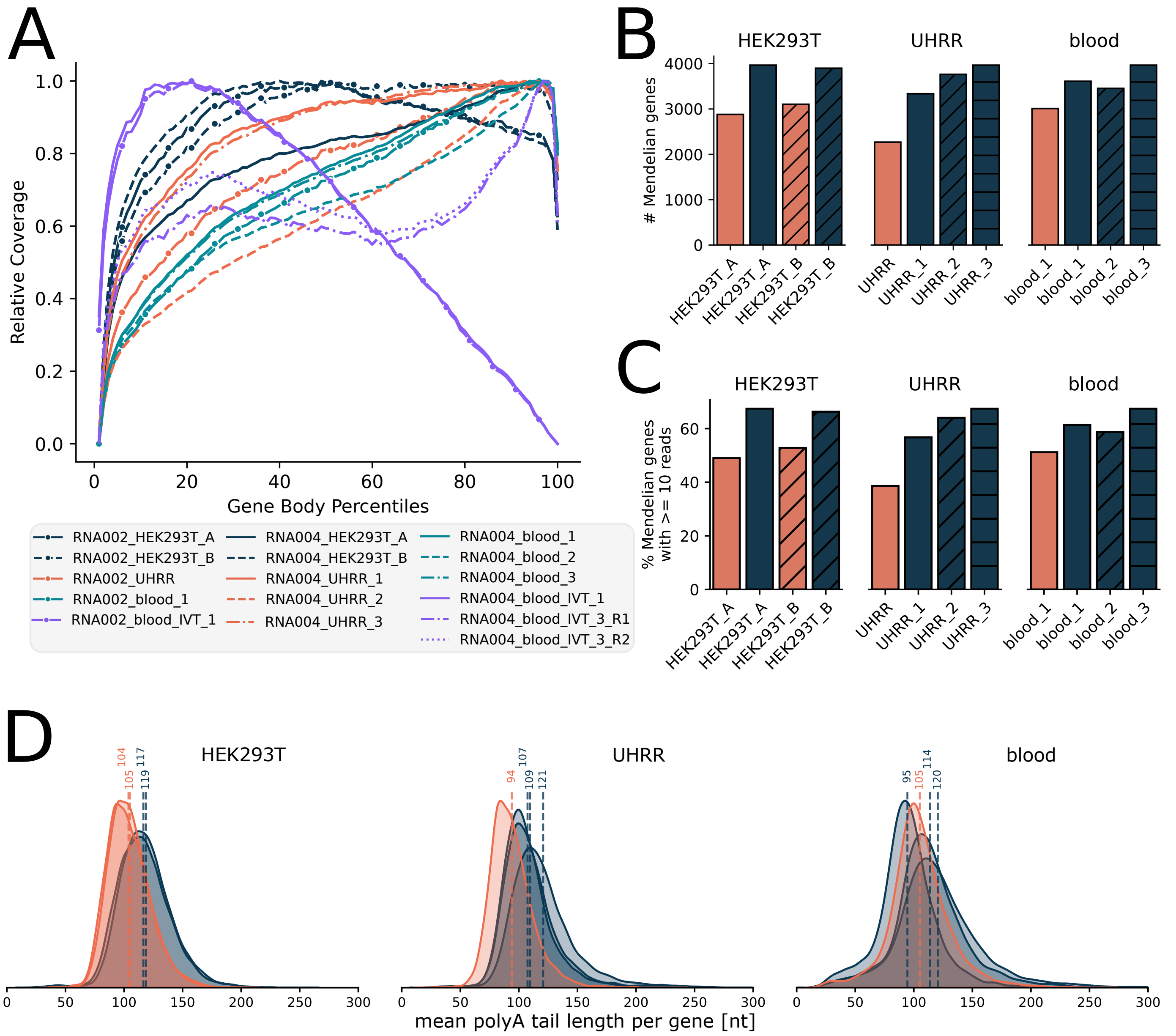


***Supplementary Figure 1.*** *(A) Transcriptome wide gene Body coverage for all samples (RSeQC, GRCh38). (B-C) Number and percentage of mendelian genes covered by HEK293T, UHRR and blood samples. (D) Density Plots of mean poly(A) lengths per gene in different samples sequenced using RNA002 and RNA004 chemistry. Samples sequenced with the RNA002 and RNA004 chemistry are shown in orange and darkblue respectively.*

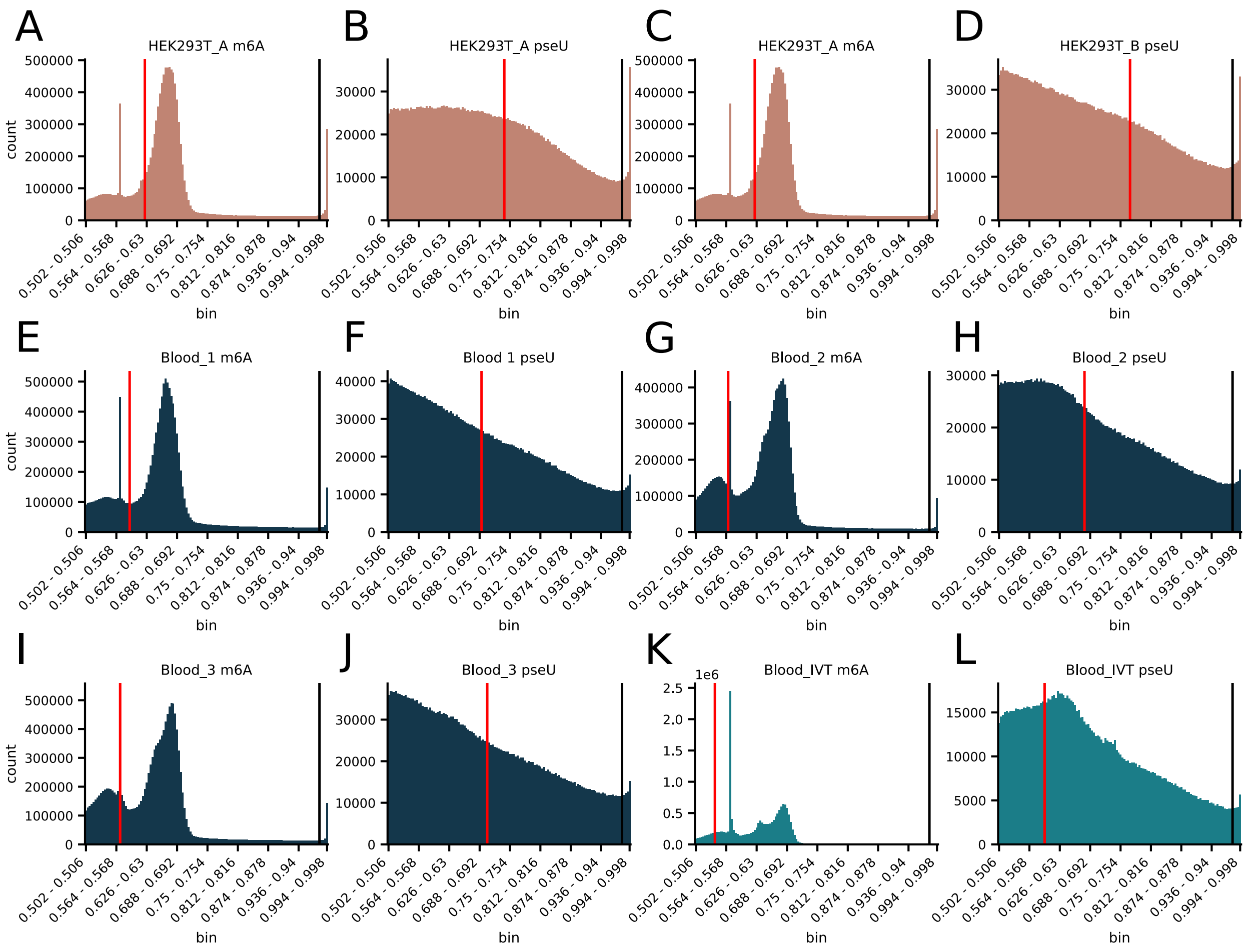


***Supplementary Figure 2.*** *Histograms of the modification probabilities of Ψ and m6A (as assigned by Dorado) for HEK293T cells (A–D) and blood replicates including IVT (E-L). The red lines indicate the filter thresholds that were selected by modkit as a default threshold (10th percentile). The black line indicates the modification threshold filter of 0.98 that was manually applied across all samples.*

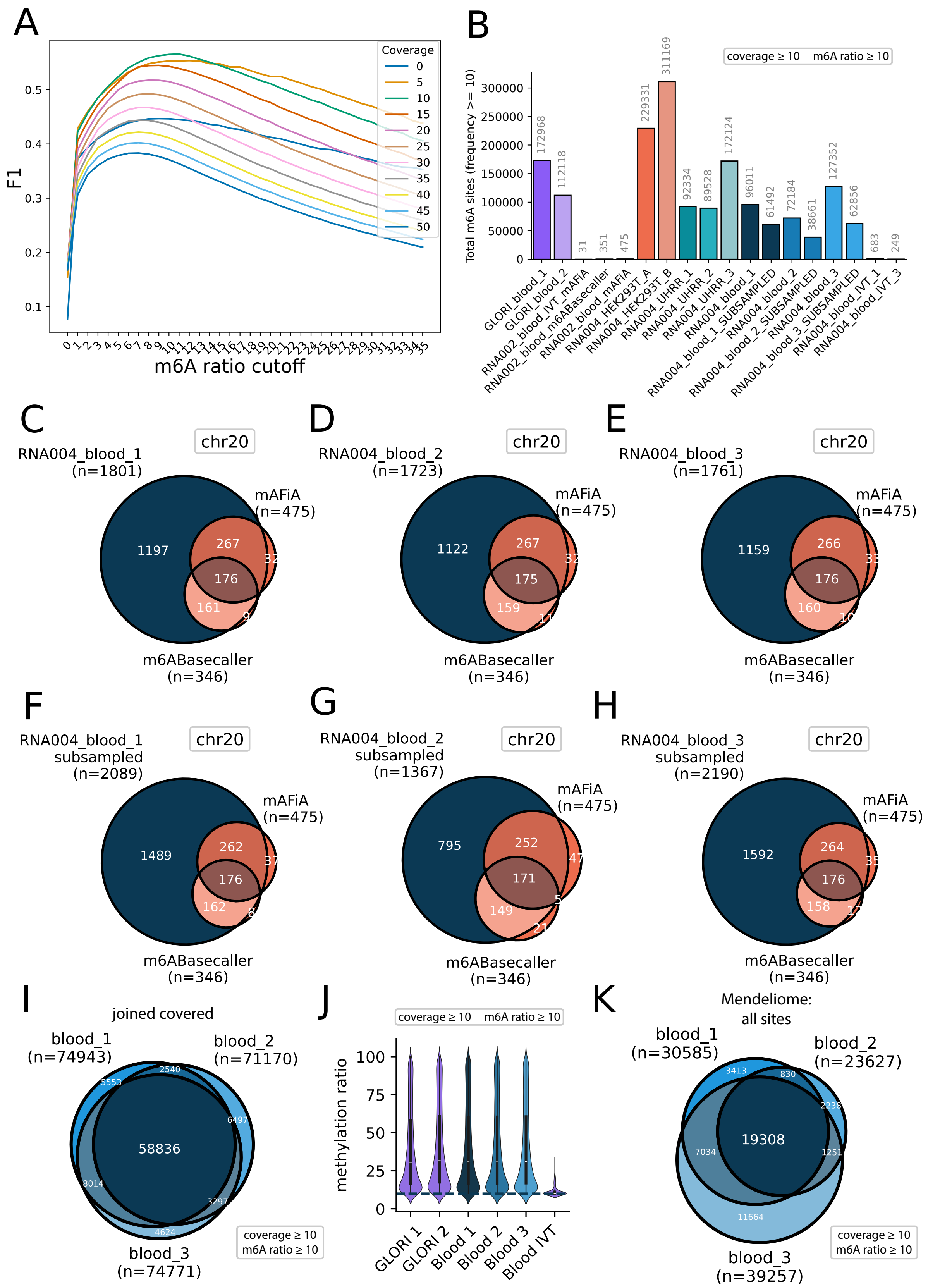


***Supplementary Figure 3.*** *(A) Plots showing F1 values at different threshold settings for coverage and methylation ratio cutoffs which were calculated by using the corresponding GLORI-seq dataset as a ground truth. (B) Total number of detected m6A sites with a ratio above 10% and minimum coverage of 10 in all samples. (C-E) Overlap between joined covered m6A sites of reads mapping to chromosome 20 of blood replicates identified by RNA002 and RNA004 using Dorado, mAFia and the m6ABasecaller. (F-H) Overlap of m6A sites between RNA002 and RNA004 m6A callers after correcting for coverage. Each RNA004 replicate shown in this intersection was downsampled to the exact readcount of the RNA002 sample. Then, set sizes were overlapped. (I) Overlap of joined covered m6A sites between blood replicates with over 10% m6A ratio and a minimum of 10 coverage. (J) Distribution of m6A ratios of joined covered sites in blood replicates with a minimum of 10 coverage and 10% m6A ratio. (K) Overlap of all m6A sites detected by DRS in blood replicates, which were found in Mendeliome genes, only sites exceeding 10 coverage and 10 m6A ratio were used for this intersection.*

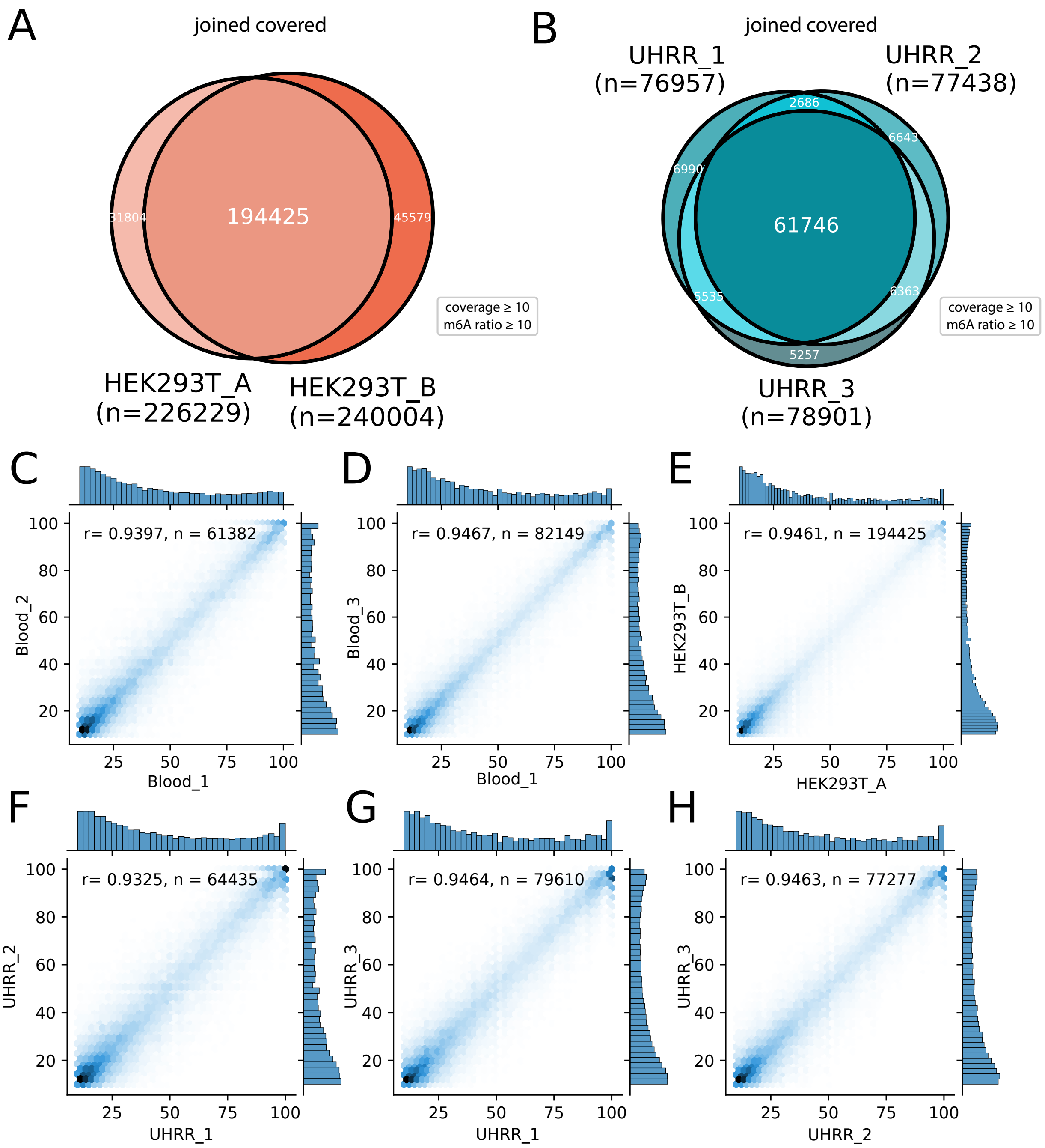


***Supplementary Figure 4.*** *(A-B) Overlap of joined covered m6A sites with a minimum of 10 coverage and 10 m6A ratio between HEK293T and UHRR replicates. (C-H) Correlation between blood, HEK293T and UHRR replicates of joined covered m6A sites detected by DRS with a minimum of 10% m6A ratio and 10 coverage. The number of joined covered sites (n) and Pearson´s correlation coefficient (r) are shown in the plots.*

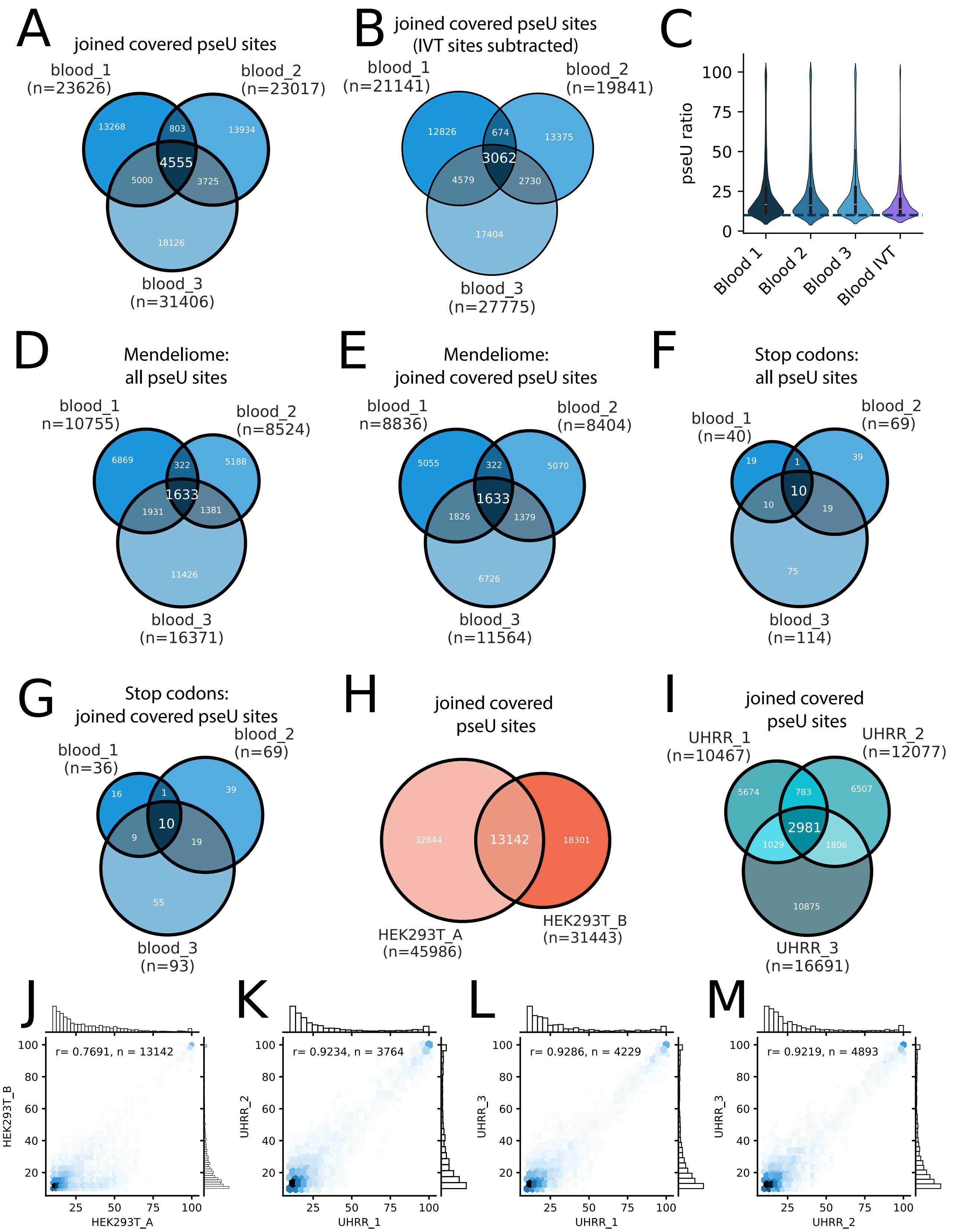


***Supplementary Figure 5.*** *(A) Overlap of joined covered Ψ sites between blood replicates with a minimum of 10 coverage and 10 modification ratio. (B) Overlap of joined covered Ψ sites with false positive IVT sites subtracted between blood replicates with a minimum of 10 coverage and 10 modification ratio. (C) Distribution of ratios of joined covered Ψ sites between blood replicates, detected by DRS. (D-G) Overlaps of either all detected Ψ sites or joined covered Ψ sites in mendeliome genes or stop codons between blood replicates. Only sites with a minimum of 10 coverage and 10 modification ratio were included. (H-I) Overlaps of joined covered Ψ sites exceeding 10 modification ratio and 10 coverage between HEK293T and UHRR replicates. (J-M) Correlation between HEK293T and UHRR replicates of joined covered Ψ sites detected by DRS with a minimum of 10% m6A ratio and 10 coverage. The number of joined covered sites (n) and Pearson´s correlation coefficient (r) are shown in the plots.*

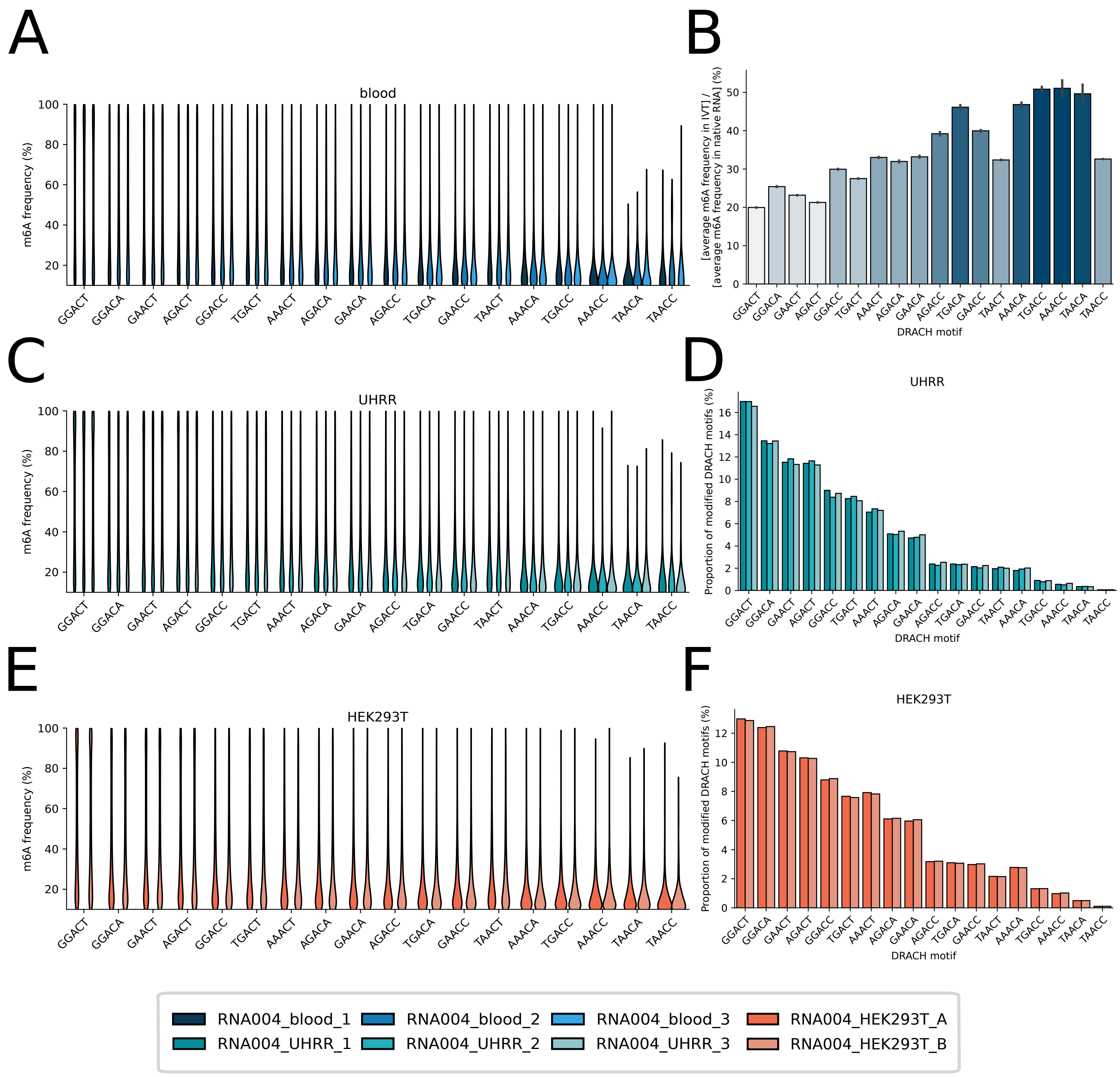


***Supplementary Figure 6.*** *(A,C,E) m6A frequencies per DRACH motif shown for all replicates in each tissue. (B) Average m6A ratio per DRACH motif in IVT shown as a ratio compared to the average m6A ratio in the respective DRACH motif detected in the native blood sample. (D and F) Proportion of modified DRACH motifs in UHRR and HEK293T replicates.*

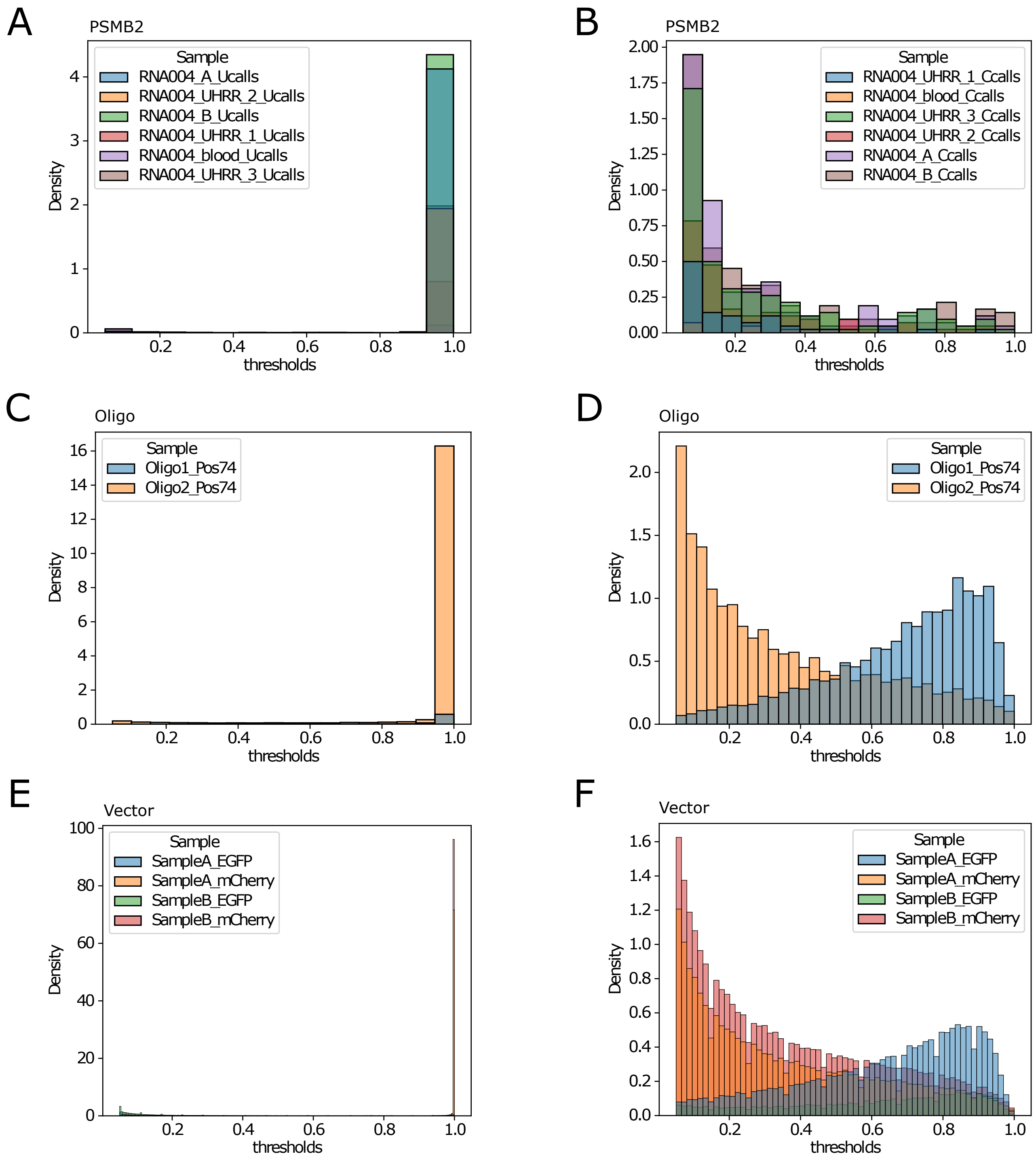


***Supplementary Figure 7.*** *Single-site Ψ modification probabilities. Plots shows Ψ modification probability for either U sites (Ucalls, left) assigned with the standard modification model or mis-base-called C sites (Ccalls, right) assigned with the modified Dorado model. (A,B) PSMB2 position chr1:35603333 for the cell line samples. (C,D) Motif-specific Ψ for the control oligos at motif1. (E,F) Ψ for the vectors EGFP and mCherry HEK293T samples A and B.*

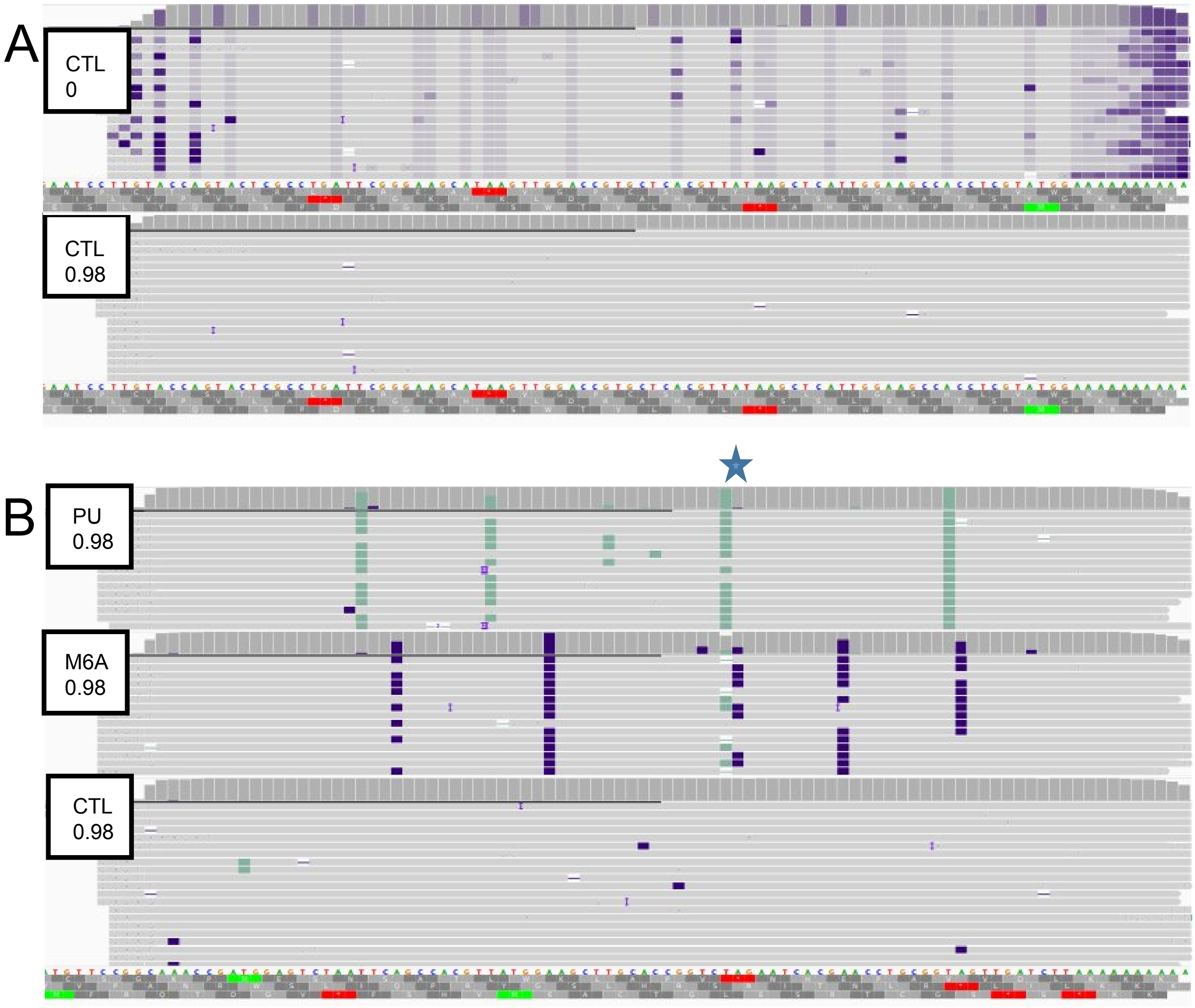


***Supplementary Figure 8.*** *Site-specific methylations of reference control oligos from ONT. (A) Upper panel: m6A-specific methylation probabilities (purple bases) per base on unmodified reference oligo 1, when a methylation threshold filter of 0 is applied, as shown in the white box. Lower panel: the exact same unmodified reference oligo 1, when a methylation threshold filter of 0.98 is applied, as shown in the white box. (B) Upper panel: pseudouridine reference oligo (pseudouridine bases are covered in light green). Middle panel: m6A reference oligo. Lower Panel: Unmodified control oligo. The basepair position of interest is marked with a green star. The middle panel (m6A control oligo) shows false positive pseudouridination calls close to the methylated base, which do not replicate at this position for the control oligo.*

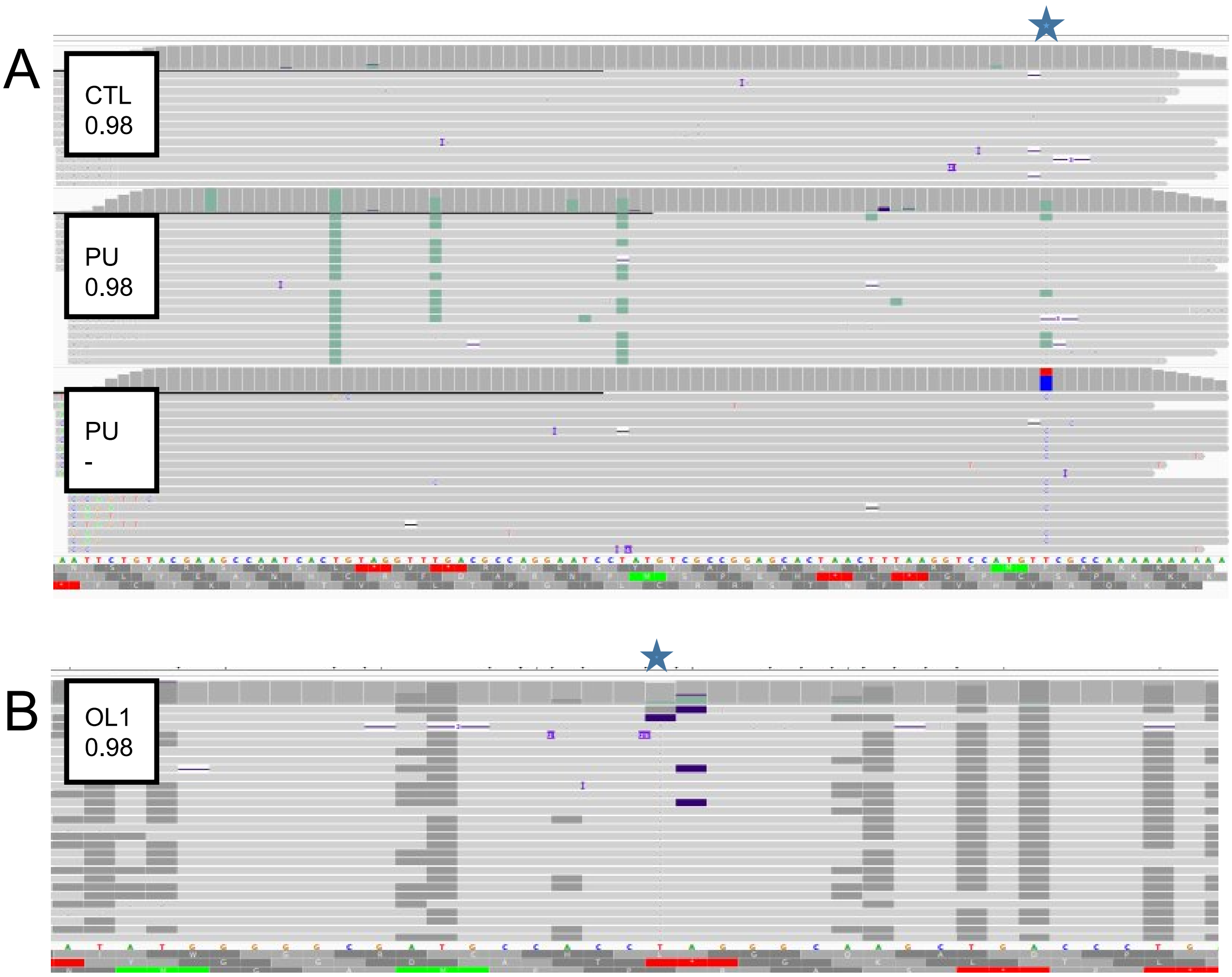


***Supplementary Figure 9.*** *U/C mismatch and site-specific methylations of reference control oligos from ONT and oligo 1 of this study. (A)The basepair position of interest is marked with a green star. Upper panel: Unmodified control oligo with a methylation threshold of 0.98. Middle panel: pseudouridine reference oligo with a methylation threshold of 0.98. Lower panel: pseudouridine reference oligo in normal view mode. At the position of interest we observe a C/U misbasecall of about 60%, that does not replicate in the control oligo. (B)Custom oligo 1 from this study. The green star marks the methylated position. We observe abberant m6A-calls directly adjacent to the methylated position.*
